# Integrating environmental gradients into breeding: application of genomic reactions norms in a perennial species

**DOI:** 10.1101/2023.11.22.568058

**Authors:** Victor Papin, Alexandre Bosc, Leopoldo Sanchez, Laurent Bouffier

**Author notes:** these authors contributed equally to this work (Leopoldo Sanchez, Laurent Bouffier). Corresponding author: Laurent Bouffier.

## Abstract

Global warming threatens the productivity of forest plantations. We propose here the integration of environmental information into a genomic evaluation scheme using individual reaction norms, to enable the quantification of resilience in forest tree improvement and conservation strategies in the coming decades. Random regression models were used to fit wood ring series, reflecting the longitudinal phenotypic plasticity of tree growth, according to various environmental gradients. The predictive performance of the models was considered to select the most relevant environmental gradient, namely a gradient derived from an ecophysiological model and combining trunk water potential and temperature. Even if the genotype ranking was preserved over most of the environmental gradient, strong genotype x environment interactions were detected in the extreme unfavorable part of the gradient, which includes environmental conditions that are very likely to increase in the future. Combining genomic information and longitudinal data allowed to predict growth in unobserved environments: considering an equivalent phenotyping effort, the cross-validation scenarios led to predictive performances ranging from 0.25 to 0.59 highlighting the importance of phenotypic data allocation. Genomic reaction norms are useful for the characterization and prediction of the function of genetic parameters and facilitate breeding in a climate change context.

## Introduction

Forest trees are keystone species in forest ecosystems supporting biological diversity and providing ecosystem services (Brockerhoff *et al*., 2017). They also produce wood, which will be a key material for meeting the challenges of the near future, thanks to its multiple uses (construction, paper, furniture, energy, chemistry) and its ability to sequester carbon for long periods of time (Ramachandran Nair *et al*., 2009; Domke *et al*., 2020). In this context, forest plantation has been expanding for several decades (FAO, 2010), with the aim of concentrating timber production and relieve pressure on natural forest. However, these benefits of forest plantation will require the adaptation of forest to a new, more challenging climate (Allen *et al*., 2010; Pawson *et al*., 2013; Payn *et al*., 2015) One of the major levers for ensuring sustainable wood productivity for forest plantations will be the deployment of trees capable of maintaining high growth rates even in extreme environments. To meet this goal, the integration of phenotypic plasticity, which is defined as the ability of a genotype to produce different phenotypes in different environmental conditions (Bradshaw, 1965), is becoming a major issue in forest tree breeding programs (Ray *et al*., 2022). A genotype is considered here as a unique genetic combination found in a single individual, or in several vegetative copies genetically identical. The challenges posed by climate change faced limited scope of traditional genetic analyses of forest trees focusing principally on phenotypic plasticity between experimental sites (Baltunis *et al*., 2010; Correia *et al*., 2010; Shalizi and Isik, 2019). These studies highlighting the existence of genotype x environment (GxE) interactions for conifer trees often consider a limited number of environments, selected so as to avoid high mortality rates. They are, therefore, not designed to be representative of the full range of environments of relevance in a context of rapid climate change. The cost and difficulty of exposing the same genotype to different environmental conditions, particularly for species difficult to propagate vegetatively, are major obstacles to the systematic evaluation of across-site plasticity in the context of tree breeding.

Phenotypic plasticity can be effectively modeled by reactions norms if repeated measurements across ages or clones are available, together with a relevant descriptor of the environment in which the phenotype was expressed (Schlichting and Pigliucci, 1998; Sanchez *et al*., 2013). A reaction norm is a representation of phenotypic values as a function of an environmental gradient. Various methods for constructing reaction norms have been developed, but the random regression model described by (Kirkpatrick and Heckman, 1989) is particularly relevant in breeding contexts. Through the integration of genetic data, this model can continuously estimate genetic parameters and breeding values according to the gradient. The gradient most frequently chosen is time (age), and this approach is frequently used in animal breeding (Jamrozik *et al*., 1997; Schaeffer, 2004; Boligon *et al*., 2012) and more rarely in plant breeding contexts (Sun *et al*., 2017; Campbell *et al*., 2018) including tree breeding (Apiolaza and Garrick, 2001; Wang *et al*., 2009). However, reaction norms along an environmental gradient have recently been modeled, as a way to meet the challenges of rapid climate change in tree breeding (Marchal *et al*., 2019; Alves *et al*., 2020).

In forest tree breeding programs, selection has historically focused on growth traits evaluated at an advanced age, they represent the cumulative reaction of the tree to the environment over a number of years (Mullin *et al*., 2011; Pâques, 2013). It is not therefore possible to trace back and identify the environmental factors contributing to the final phenotype, as environment can be considered only in a global manner over the whole period. However, yearly growth increments can be correlated with well-characterized environments (Martinez-Meier *et al*., 2008; Zas *et al*., 2020). This can be achieved with the use of wood ring series, which define the annual radial growth of each individual in temperate climates. Indeed, the cambial activity of trees depends strongly on environmental conditions, particularly temperature and water availability (Schweingruber, 2007). The variability of annual ring width and wood density characterizes the plastic response of trees to changing environmental conditions. It has been shown to have genetic determinism (Sánchez-Vargas *et al*., 2007; Dalla-Salda *et al*., 2009) and could be used as a proxy for the potential reaction of trees to changes in environmental conditions. The analysis of these repeated phenotypes therefore provides an ideal longitudinal dataset for studying phenotypic plasticity at individual level (Marchal *et al*., 2019). Such analyses can be explanatory in nature, seeking to identify the optimal combination of environmental factors making a significant contribution to annual growth, but they can also be predictive, with the development of functional models for inferring growth in unobserved environments.

The integration of molecular markers into genetic evaluations provides not only more accurate estimates of genetic parameters, but also opportunities to implement genomic selection (GS) approaches (R2D2 Consortium *et al*., 2021). In forest trees, such approaches pave the way for the early selection of important traits, such as wood traits, that would otherwise be evaluable only after many years of cumulative growth. GS is also particularly valuable in tree breeding, as it allows the integration of traits that are costly and complex to measure (Grattapaglia and Resende, 2011). In many species, the gains provided by the use of genomic data have tended to eclipse the interest in longitudinal data (Oliveira *et al*., 2019). However, these two approaches are not antagonistic and their beneficial effects can be combined (Rutkoski *et al*., 2016; Sun *et al*., 2017). Genomic reaction norms are rarely used (Ly *et al*., 2018), but are potentially of great value in this context, as they allow prediction of growth in as yet unobserved environments, thus decreasing the complex and costly evaluation procedures associated with experimentation and phenotyping under different environmental conditions.

We propose here an integration of environmental information into genetic evaluations, using reaction norms in the context of forest tree breeding. A random regression model based on annual ring growth data for maritime pine (*Pinus pinaster* Ait.) and including genomic data was used to fit individual-level reaction norms. The genetic components of these norms were described and the implications of their use in the context of breeding were further investigated with respect to a classical analysis targeting final radial growth. Finally, we investigated the model’s ability to predict growth in unobserved environments considering realistic phenotyping conditions for the maritime pine breeding program in a GS context. To our knowledge, this is the first study in a tree breeding context to use a random regression model to combine environmental gradient and genomic information.

## Materials and Methods

### Plant material

A maritime pine trial was established at two sites in 1997: Site 1 (Cestas: Lat 44.74, Lng −0.68) and Site 2 (Escource: Lat 44.16, Lng −1.03). Soil characterization revealed greater soil fertility (+16.8 g organic matter/kg of soil) and a shallower water table (mean difference of +6 m) at Site 1 than at Site 2. Climatic measurements showed that there was more rainfall at Site 2 (mean of +15% for total annual rainfall), whereas temperatures were equivalent at the two sites (supplementary Table S1). A total of 202 genetic units (35 trees per genetic unit) were studied on both sites. They were planted in a complete block design with single-tree plots (1,250 trees/ha) and consisted of 196 half-sib families obtained from crosses between identified seed parents and two pollen mixtures of identified donors, plus six check lots. Thinning operations were performed at both sites in 2012 and exclusively at Site 1 in 2017, when the trees were 16 and 21 years old, respectively. A subsample (POP) of 25 half-sib families, with 13 individuals per family and per site, was selected as representative of the variability of growth (total of 650 individuals). In this maritime pine experimental context, each genotype is represented by a single individual, so that the notions of “genotype”, “individual” and “tree” are considered equivalent in our study.

### Genetic characterization of POP

Genomic DNA was extracted from needles collected from POP, to which we added 186 randomly selected duplicates for repeatability estimates. The concentration and quality of DNA for each sample were determined with a NanoDrop spectrophotometer (NanoDrop Technologies, Wilmington, DE, USA). Genotyping was performed by Thermo Fisher Scientific (Thermo Fisher Scientific, Santa Clara, CA, USA) with the 4TREE Axiom 50K single nucleotide polymorphism (SNP) multi- species array (Guilbaud *et al*., 2020). The preliminary filters recommended by Thermo Fisher Scientific were applied to the genotyping results, at the sample (DishQC ≥ 0.4, CallRate ≥ 90) and SNP (CallRate ≥ 95, fld-cutoff ≥ 3.2, het-so- cutoff: ≥ −0.1) levels. In addition, sequential filtering was applied, with the removal, in the following order, of SNPs with less than 85% repeatability, SNPs with more than 5% Mendelian segregation errors and SNPs with a minor allele frequency (MAF) below 1%. A genomic relationship matrix (G) was calculated with the VanRaden formula (VanRaden, 2008) using the AGHmatrix package (Amadeu *et al*., 2016) in R 4.2.2 environment (R Core Team, 2022):

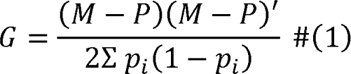

where the *M* matrix (*n*: number of individuals *x m*: number of markers) contains marker information coded as −1 for one of the homozygotes, 0 for heterozygotes and 1 for the other homozygotes; and the *P* matrix (*n* × *p*) contains allele frequencies expressed as 2(*p_*i*_* − 0.5), where *p_i_* is the frequency of the second allele at locus *i* for all individuals.

In addition, pedigree recovery was performed for each tree from POP, with a subset of 161 SNPs used to infer the identities of the parents (25 seed parents and 85 pollen parents) and grandparents (69 initial progenitors from the base population of the breeding program) (supplementary Method. S1). The most complete version of the pedigree was used to compute an additive relationship matrix A for further analyses.

### Phenotypic data

Circumference measurements were performed on all the trees in the trial in 2004, 2008, 2012 and 2018, at the ages of 8, 12, 16 and 22 years, respectively. In addition, cores were removed from the trees of POP in December 2019, at breast height, along the same north-south direction for each tree. These cores were cut into 2-mm-thick radial strips for X-ray analysis (Polge, 1966) to obtain wood density profiles (Fig. 1). The limits between the different rings were identified with Windendro software (Guay *et al*., 1992) and validated by visual examination. The area of ring (*RA_raw_y__*) was calculated at individual level as follows:

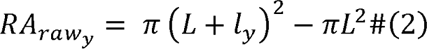

**Figure 1:**
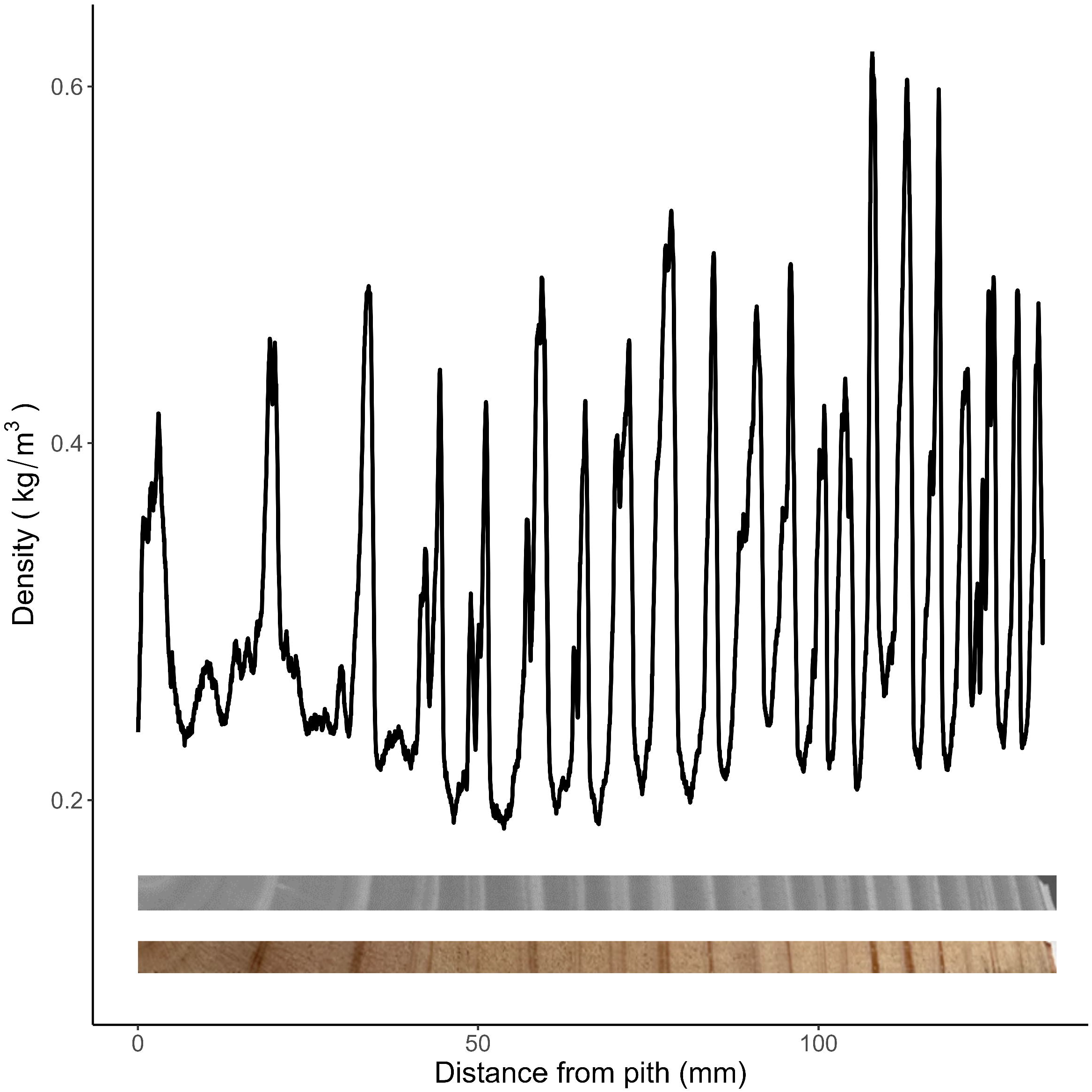
From wood increment core to wood density profile. From the bottom to the top: the wood increment core picture from one tree, its corresponding radiography, its wood density profile (black line) from pith (position: 0mm) to the bark obtained after processing. Sudden and high drops in wood density mark the end of annual growth and were used to fit each ring limitations.

where *L* is the sum of the ring widths from the pith to ring *y* (ring *y* excluded) and *l_y_* is the width of ring *y*. *RA_raw_* values are a good proxy for biomass produced each year independently of tree age, in contrast to ring widths which tend to decrease progressively over the years due to radial growth of the tree.

We chose to study the 2005-2019 period (15 successive years) here because rings for this period were available for at least 99% of POP and this period excludes the juvenile phase of the trees (supplementary Table S2 and Fig. S1). Using the circumference measurements, *RA_raw_* values were spatially corrected for each site with spline functions (via the BreedR R package; Muñoz and Sanchez, 2020) and named RA (adjusted ring area). A complete phenotyping series for an individual is thus composed of 15 RA values.

#### Characterization of the environment during ring growth

The environmental conditions associated to each ring were characterized with two classes of environmental indices, which depend on both year and site variables. The first class focused on a purely climatic description, with two versions (*DM* and *DM′*) of the de Martonne aridity index (de Martonne, 1926), whereas the second provided a finer description of the environmental conditions with two indices (*GP* and *GP′*), extracted from an ecophysiological model combining climatic, silvicultural and soil data (Moreaux *et al*., 2020).

The de Martonne aridity index was calculated for each ring formed in year *y* at site *Z* with:

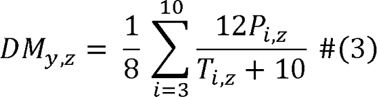

Where *P_i,z_* is the amount of precipitation (in mm) and *T_i,z_* is the mean air temperature (in °C), for month *i* in site *z* Only the 8 months from March (*i* = 3) to October (*i* = 10) were included here as we considered, as a first approximation, that climatic conditions outside the growth period of maritime pine has no impact on annual RA. In addition, we considered a modified version of the de Martonne index (*DM′*) based on a 30-day sliding window average (instead of calendar months) and considering the impact of the climate of year (*y* − 1) on environmental conditions in year *y* (inspired by Botzan *et al.*, 1998, supplementary Method S2A).

The environmental indices of the second class derived from the ecophysiological model GO+ 3.0 (Moreaux *et al*., 2020) based on climatic data, silvicultural parameters, soil water properties, soil fertility and reference values for maritime pine growing in the Landes massif (supplementary Method S2B). The growth potential index (*GP*) was calculated for each ring, based on mean trunk water potential and temperature estimated daily by the GO+ model (supplementary Method S2C). Similarly, to the de Martonne aridity indices, a second index *GP′* was used to consider a sliding window of 10 days over the course of a year and to take into account the impact of previous year.

### Genetic analysis of radial growth

#### Univariate model

Radial growth analysis for POP would classically be based on the most recent available circumference measurement (here for 2018, when the trees were 22 years old, denoted Cir22) and the following univariate model:

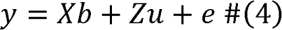

Where *y* is the vector of Cir22; *b* and *u* are the vectors of solutions for fixed site and random genetic additive effects, respectively; *e* denotes the residuals. *X* and *z* are the corresponding incidence matrices of dimensions *n x* 1 and *n x n,* respectively, where *n* is the number of individuals. We assumed that 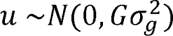 and 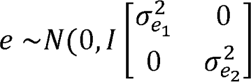), with *G* the genomic relationship matrix described above, 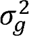 the additive genetic variance, 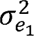 and 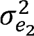 the residual variances for Site 1 and Site 2 respectively. The genomic estimated breeding values (GEBV) for Cir22 will be denoted GEBV_Cir22_.

#### Random regression model (RRM)

Individual RA series for POP were modeled as a function of the environmental gradient, using an RRM implemented in Wombat software (Meyer, 2007). The environmental gradients associated with the four indices previously described were modeled independently according to the RRM formulation. Regardless of the environmental index used, the joint analysis of the two sites and year series provided an overall environmental gradient of 30 levels (15 environmental levels per site). Legendre polynomials were used as the base functions (Kirkpatrick *et al*., 1990) for the following RRM (Mrode and Thompson, 2005):

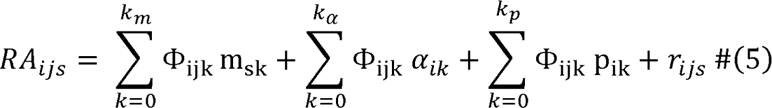

where *RA_ijs_* is the ring area of individual *i* for environmental level *j* at site; *m_sk_* is the *k*^th^ fixed regression coefficient used to model the average trajectory at site *S*; *α_ik_* and *p_ki_* are the *k*^th^ random regression coefficients for the genetic additive and permanent environmental effects, respectively, of individual *i*, the latter effect representing the similarity between repeated records for the same individual of environmental and non-additive genetic origin; Φ_ijk_ is the *k*^th^ Legendre polynomial for the RA of individual *i* at environmental level; *j*; *k_m_*, *k_α_*, *k_p_* are the order of polynomials for mean trajectory, genetic additive and permanent environmental effects, respectively; and *r_ijs_* is a random residual. Based on the results of a preliminary analysis (supplementary Fig. S2), we decided to model second-order trajectories by setting *k_m_* = *k_α_* = *k_p_* = 2.

The equivalent matrix notation for this model is (Mrode and Thompson, 2005):

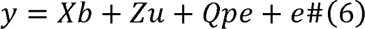

where *y* is the vector of RA over the environmental levels; *b* is the vector of solutions for site fixed effect; *u* and *pe* are the vectors of the individual genetic additive and permanent environmental random regression coefficients, respectively; *e* denotes the residuals. *X,Y*, and *Q* are the corresponding incidence matrices; *Z* and *Q* being of rank (*n_j_ x n_k_*) where *n_j_* is the number of environmental levels, and *n_k_* is *k_α_* times the number of genotypes with records. For genomic-based RRM, it is assumed that *u ~ N*(0,*G* ⊗ Ω), *Pe* ~ *N* (0, *I* Ω *P*) and *e* ~ *N* (0, *I* ⊗ *D*), where ⊗ denotes the Kronecker product, *G* the relationship matrix described above, *Ω* and *P* the covariance matrices for the RR coefficients for the genetic additive and permanent environmental effects, respectively, and *D* is a diagonal matrix of heterogeneous residuals for each environmental level. For pedigree-based RRM, *G* is replaced by *A*.

With a second-order model (*k_m_* = *k_α_* = *k_p_* = 2), the RRM estimates three genetic coefficients per individual. From these, individual GEBV were then obtained at all environmental levels as a trajectory, following the formulation of (Mrode and Thompson, 2005). GEBV estimated with an RRM integrating all available phenotypic data and solved at each environmental level are denoted GEBV_ref_. The individual trajectories of GEBV_ref_ as a function of the environmental gradient were clustered with a K-means approach extended to longitudinal data (implemented in the kml R package; Genolini *et al*., 2015). The Calinski-Harabatz criterion was used to define the best number of clusters (ranging from 2 to 7).

### Genomic selection

#### Cross-validation (CV) scenarios

The predictive performance of the RRM was assessed over two CV scenarios (Fig. 2). First, the reference scenario, denoted **CV-A**, where the training set (T_set_) included the complete phenotyping series for 50% of the individuals (randomly selected within sites and families), whilst the remaining 50% of individuals constituting the validation set (V_set_). Second, the CV-B scenario explored the possibility of retaining the same amount of phenotypic information as for the CV- A (i.e. 50% of total phenotypic data) but distributed differently over the individuals. Scenario **CV-B** mimicked the use of a high-throughput phenotyping tool for quick estimation of the last five RA which, in a context of global warming, would typically correspond to unfavorable years. The T_set_ for **CV-B** included complete phenotypic series (i.e. 15 phenotypic records per individual) for 25% of individuals and only five phenotypic records for the remaining individuals (75% of individuals). We selected the five environmental levels within the eight most unfavorable ones by applying a Kennard-Stone algorithm (Kennard and Stone, 1969) via the prospectr R package (Stevens and Ramirez-Lopez, 2022).

**Figure 2:**
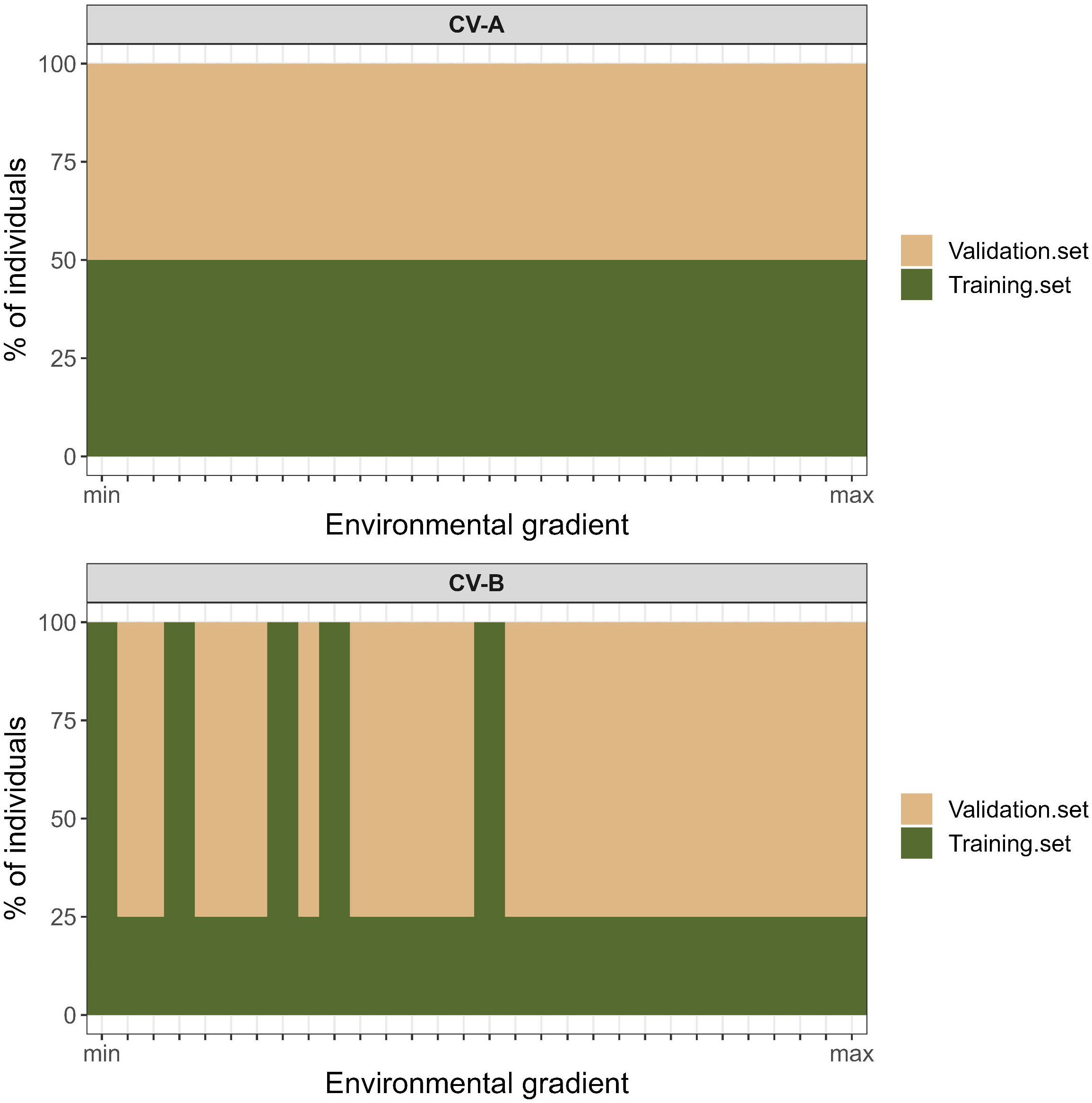
Cross-validation scenarios CV-A, CV-B, CV-C and CV-D performed with a RRM according to the *GP′* index. All scenarios include the same amount of phenotypic information in the training set (i.e. 50% of total phenotypic data); only the distribution of this information across individuals and environmental levels differ. All families contributed equally to the training set.

For each CV scenario, the predictive performance of the RRM was estimated as the Pearson correlation coefficient between predicted (GEBV_pred_) and observed RA in V_set_ over the whole environmental gradient and based on 10 independent repetitions. Such performance estimator was used as a criterion for assessing modeling quality (Ly *et al*., 2018; Arnal *et al*., 2019; Momen *et al*., 2019).

#### Genetic gains

The predictive performance of the RRM for genetic gains in our reference scenario CV-A was assessed over each of the environmental levels. The assessment consisted of calculating the differences in genetic gain between a selection based on GEBV_pred_ obtained in V_set_ and the corresponding maximum that would have been obtained with the same selection intensity based on GEBV_ref_. For this, at each environmental level, the top 5% of individuals selected according to GEBV_pred_ were identified and their corresponding GEBV_ref_ (obtained with all the phenotypic information) used to calculate the true genetic gain (GG_true_) as the GEBV_ref_ average of the selected individuals. This amount was compared for the corresponding environmental level to the maximum gain (GG_max_), which was calculated as the GEBV_ref_ average of the top 5%. Finally, GG_true_ and GG_max_ were centered and reduced to ensure comparability between environmental levels. Any difference between GG_true_ and GG_max_ would indicate a decrease in the correlation between GEBV_pred_ and GEBV_ref_ for the selected percentage.

## Results

### Size and genetic characterization of POP

After phenotype curation (9 wood-density profiles excluded) and genotyping quality control (13 genotypes excluded), POP was finally composed of 628 trees (303 from Site 1 and 325 from Site 2).

Pedigree recovery on POP validated 93% of the pedigree seed parents (monoicous individuals acting as mothers) and allowed the correction of 5%. The remaining 2% of the pedigree seed parents was classified as unknown, as no candidate parent could be validated. Pollen parents (acting as fathers) were successfully recovered for 65% of the individuals. Note that the original design of the study was based on crosses with a mixture of pollen donors, resulting in the fathers initially being unknown in the pedigree. Finally, based on the curated pedigree, a status number (_s_; Lindgren *et al*., 1996) of 21 was obtained for POP, suggesting a high level of relatedness between the families studied.

The genotyping of POP resulted in the characterization of the 628 individuals over 3,832 SNPs, with a repeatability of 97% and a total missing data rate of 1%. Genomic relationship coefficients (*g_xy_*) estimated in were consistent with the additive relationship coefficients (*g_xy_*) calculated in (Fig. 3). The *a_xy_* values were discrete, whereas the *g_xy_* values were normally distributed for each level of relatedness. Note that, for most additive coefficient levels, the normal distribution has a long upward-sloping tail (revealing some rare cases of unrecorded relatedness), and a mean slightly below the theoretical value, the latter being represented by the gray line in Fig. 3.

**Figure 3:**
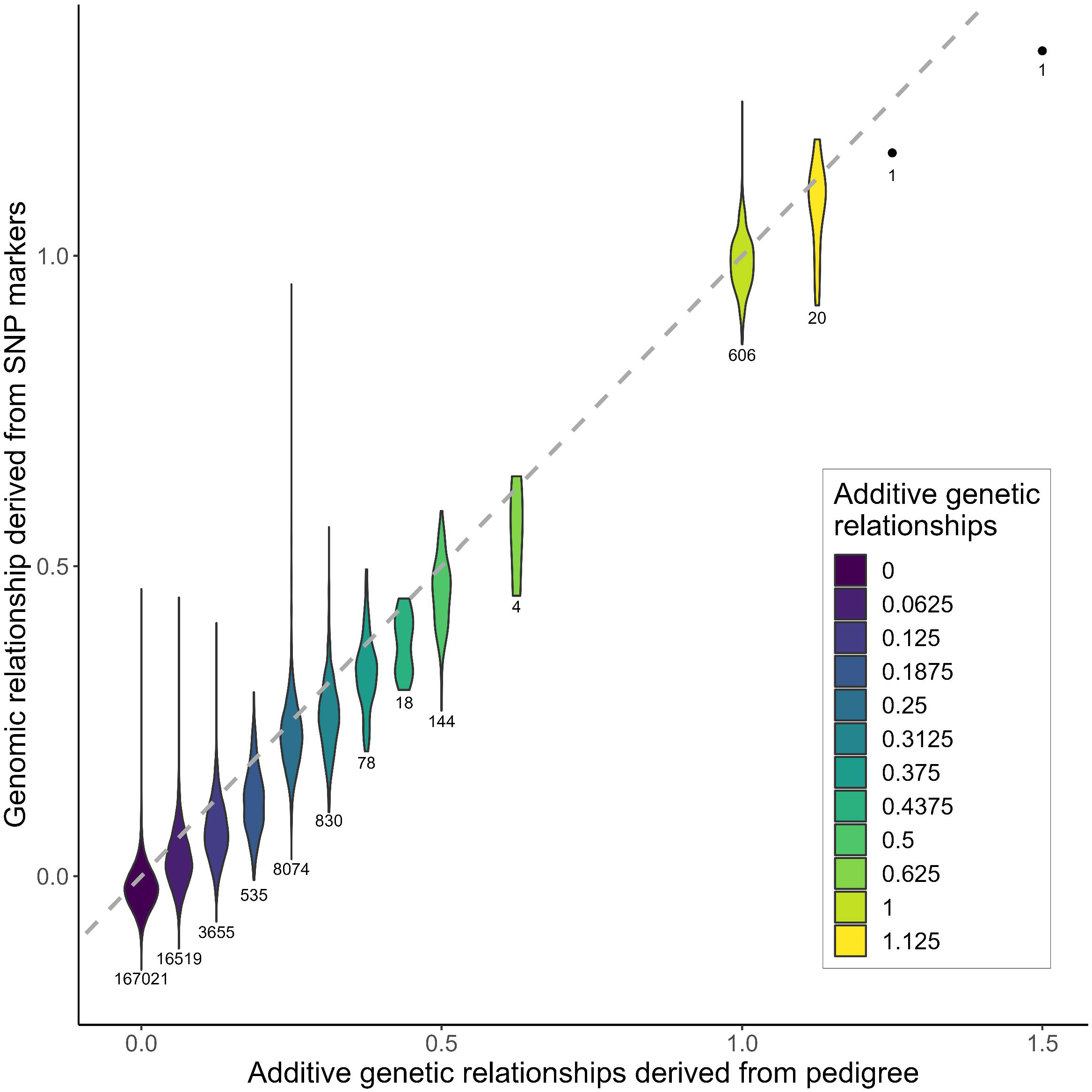
Comparison between additive genetic relationships derived from pedigree and genomic relationships derived from SNP markers for individuals of POP. *For each value of the discrete scale taken by the additive genetic relationships, the corresponding violin plot represents the continuous distribution of genomic relationships. Numbers below each violin plot denote the number of relationship included in the corresponding violin plot. Grey line is the bisector passing through the origin of the graph. The two highest relationships derived from pedigree (1.25 and 1.5) are unique and so represented by single points instead of violin plots*.

### Quality of model fit

The predictive performance (estimated with CV-A) was used as a criterion for assessing the quality of RRM (Fig. 4). Mean predictive performances were poor, with correlation coefficients ranging from 0.19 to 0.25. Predictive performance was slightly better (+0.04 better, on average) for genomic-based RRM than for pedigree-based RRM, except for RRM based on the *DM* environmental index (equivalent mean predictive performance of 0.21). The best predictive performances were obtained for genomic-based RRM with the *DM′* (0.24) and *GP′* (0.25) indices. The optimization of environmental indices improved slightly RRM predictive ability by 16% and 3% relative to the initial *DM* and *GP* indices, respectively. Finally, the genomic-based RRM using the *GP′* index was selected for the analyses described below, due to its best predictive performance (0.25) for GS.

**Figure 4:**
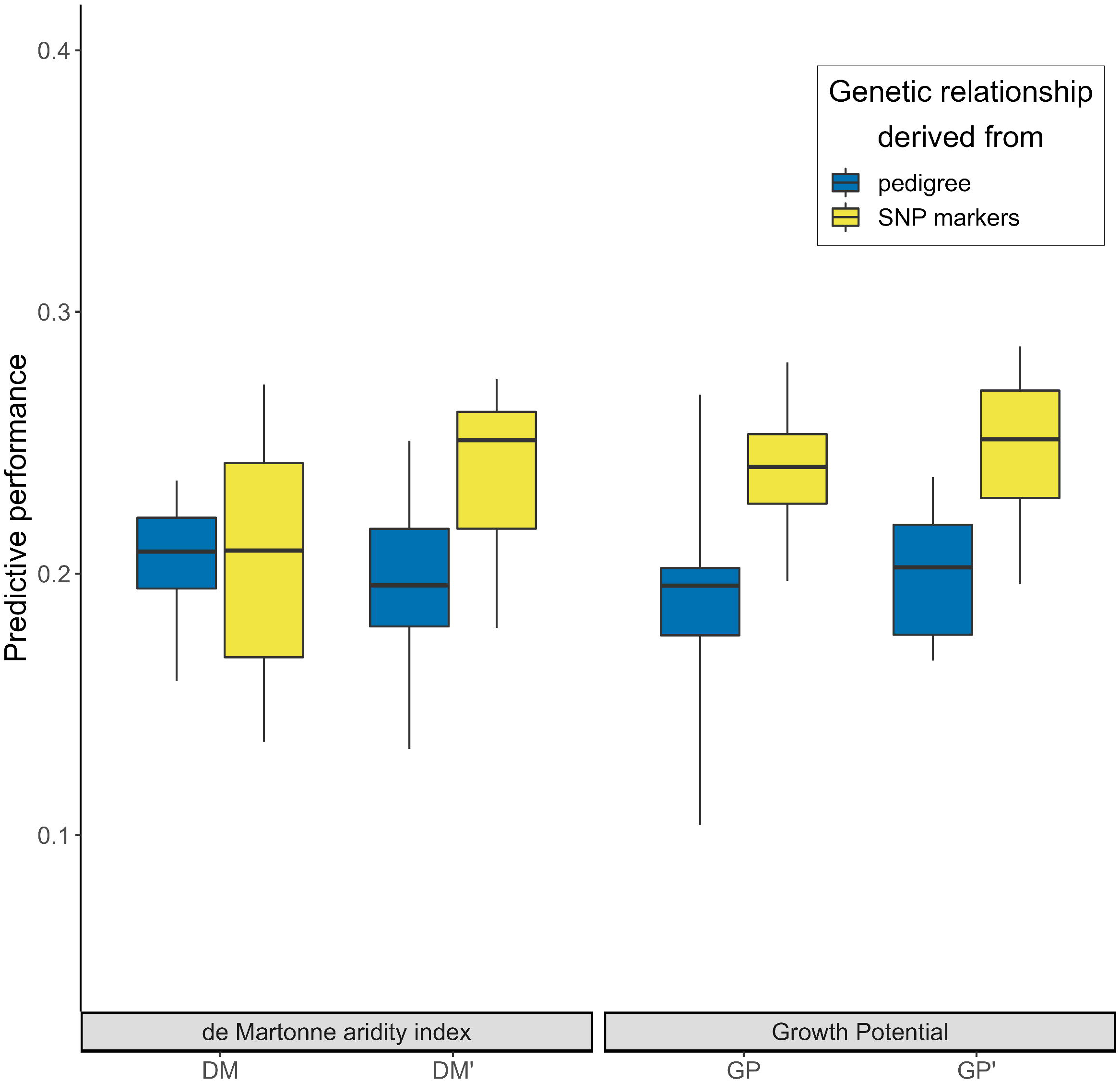
Predictive performance of the RRM according to the environmental gradient and the genetic information used. Boxplots indicates the Pearson correlation coefficient between observed and predicted RA values over the whole environmental gradient for 10 repetitions of the CV-A scenario. Boxplots are blue when the RRM implemented integrated additive genetic relationship derived from pedigree while boxplots are yellow when the RRM implemented integrated relationships derived from SNP markers. For each kind of genetic information, RRM were run independently with each of the four environmental gradients, respectively derived from, *DM*, *DM′*, *GP* and *GP′* indices.

### Individual reactions norms estimated by genomic-based RRM

Reordering longitudinal data by the annual environmental index, which characterizes the conditions of ring formation, instead of the ordinal year greatly modified the shape of the mean RA curve in a more easily interpretable way (Fig. 5). When expressed as a function of the environmental index *GP′*, RA increases significantly. The lowest *GP′* values are associated with the most unfavorable environmental conditions for growth, whereas the highest values are associated with the most favorable conditions for growth. This pattern suggests plasticity at the population level, but hides individual behaviors, which may deviate from this central trajectory.

**Figure 5:**
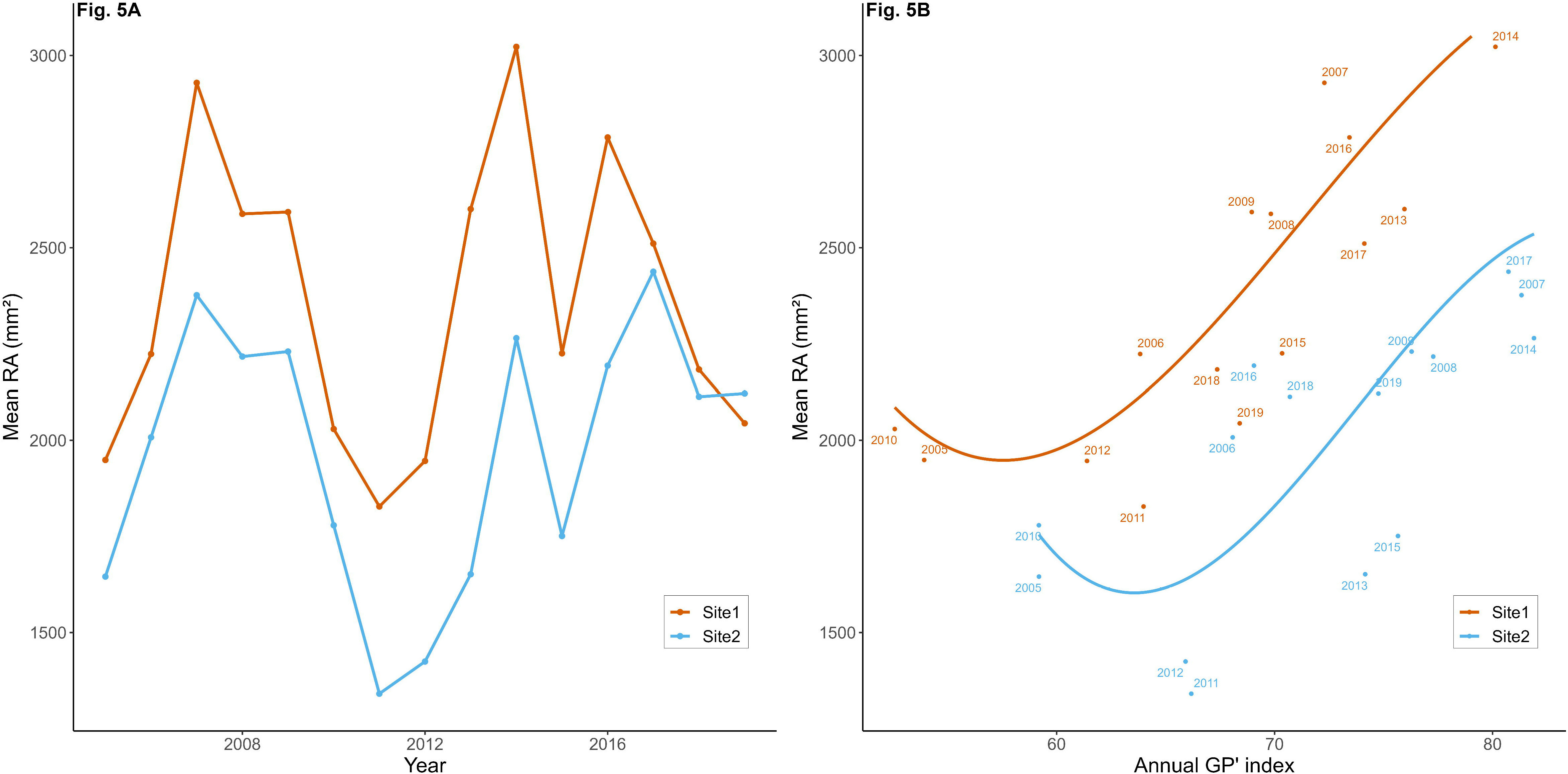
Evolution of mean RA according to the years for each site (Fig. 5A) or according to the GP′ index (Fig. 5B). Figure 5.A presents mean phenotypic trajectories of RA and Figure 5.B presents mean trajectories adjusted by the RRM for each site. Both trajectories are the result of the same model. The significance of the slope parameter for each trajectory in Figure 5.B was assessed with Student’s t-test (p-value<0.01)

Random individual deviations from the mean trajectories due to additive genetic effects are represented in Fig. 6 and were solved over the environmental gradient of *GP′* (GEBV_ref_). For most genotypes, GEBV_ref_ showed a dependence on *GP′*, highlighting the existence of plasticity for RA. These different behaviors can be characterized thanks to five clusters, depicted in different colors (Fig. 6). Each individual reaction norm can be characterized by its mean GEBV_ref_ and its slope defined with two coefficients (quadratic and linear regression coefficients as second-order trajectories were modelled). Fifty-seven percent of individual reaction norms were assigned to clusters A and B (Fig. 6) and were characterized by shallow slopes (supplementary Fig. S3) and mean GEBV_ref_ close to 0 (supplementary Fig. S4). This does not mean that the phenotypic trajectory of these individuals is flat. Instead, it indicates that they have trajectories indistinguishable from the mean trajectory and that this indistinguishability, by its additive genetic origin, would not give extra plasticity to the offspring. The highest and lowest mean GEBV_ref_ were those obtained for individuals from clusters C-E (26%) and cluster D (17%), respectively. These clusters also display reaction norms with the strongest positive and negative slopes (for clusters C-E and D, respectively), leading to a greater range of variation in individual genetic values in favorable than in unfavorable environmental levels. Mean GEBV_ref_ is, thus, strongly correlated with slope regression coefficients (0.52 and 0.95 with quadratic and linear regression coefficients, respectively). There appear to be few intersections between reaction norms, corresponding to changes in individual ranks across environmental levels, over most of the environmental gradient.

**Figure 6:**
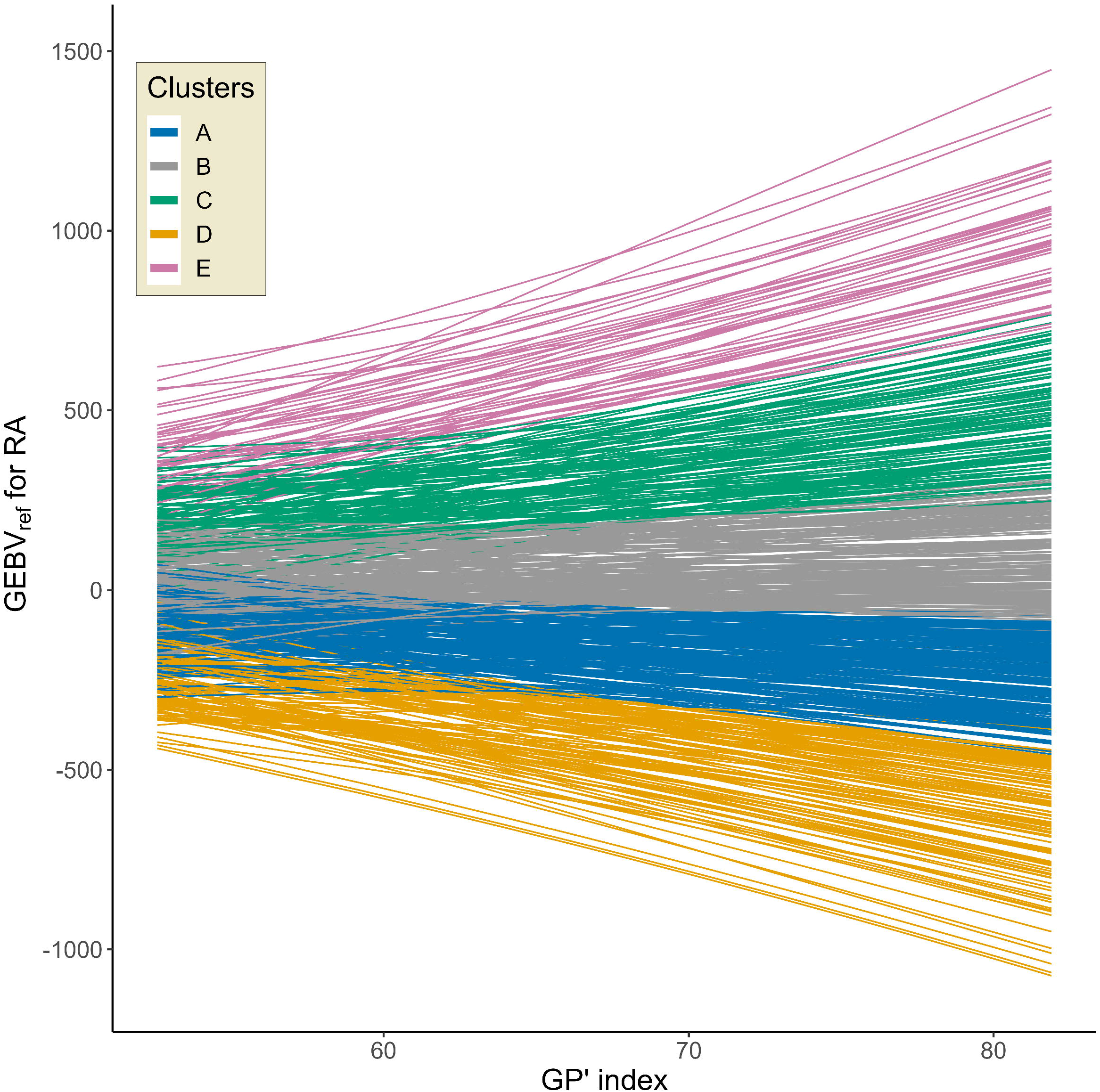
Individual trajectories of GEBV_ref_ associated to RA according to the GP′ index. Trajectories correspond to the genetic component of the reaction norms estimated by the genomic based RRM. Trajectories were divided in 5 clusters (A to E) with the following proportions: A-29.6%, B-27.3%, C-18.6%, D-17.2%, E-7.3%.

However, large overlaps occur in the part of the gradient corresponding to unfavorable environmental levels. Moreover, despite the overall similarity of trajectories within the same cluster, overlaps between norms were detected within clusters, mostly in the unfavorable part of the gradient (supplementary Fig. S5).

Correlations were calculated between the GEBV obtained with the final-point univariate model (GEBV_FP_) and those obtained with the RRM across the successive environmental levels *j* (GEBV_ref_j_). For GP′ environmental levels over 59, correlations remained almost constant ranging from 0.77 to 0.78, highlighting the consistency between GEBV_ref_j_ and GEBV_FP_ at most environmental levels (supplementary Fig. S6). However, the correlation was clearly weaker for the most unfavorable environmental levels (GP′<59), with the coefficient falling to a minimum of 0.74. These lower values highlight differences in the behavior of some genotypes (and, therefore, genotype ranking) at these environmental levels.

### Genomic selection scenarios and cross-validation

#### Genetic gain over the environmental gradient

The overall predictive performance of the genomic-based RRM (using the GP′ environmental index) estimated with the CV-A scenario was 0.25 (Fig. 4). Breeding efficiency, based on predicted values, was assessed by calculating genetic gains for different environmental levels (Fig. 7). GG_max_ increased until the environmental value of 62, above which it reached a plateau with maximum value of 2.35. The differences between GG_max_ and GG_true_ was minimal for GP′ environmental level 0.64, increasing to a maximum at the most unfavorable (0.73) and most favorable (0.72) environmental levels. The relatively poor predictive performance of the RRM (0.25) necessarily led to a significant loss of genetic gain (no overlap between GG_max_ and GG_true_ boxplots). Nevertheless, depending on the environmental level, GG_true_ accounted for 68% to 73% of GG_max_. GG_true_ was always significantly different from 0 (*p − value_t-test_* < 0.001), indicating a certain efficiency of selection based on predicted values, even in the most extreme environmental levels.

**Figure 7:**
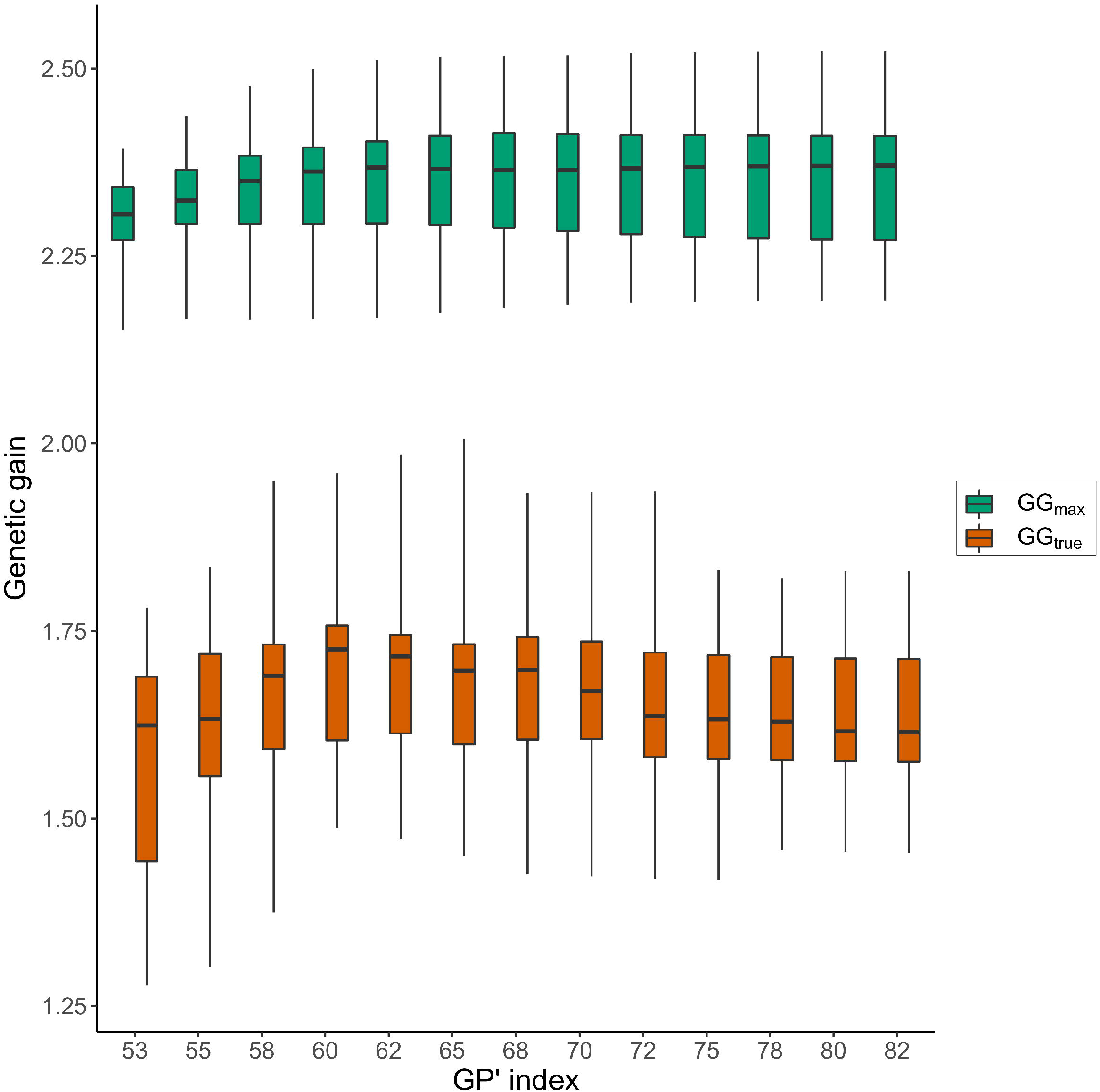
Maximum genetic gain (GG_max_) and true genetic gain (GG_true_) according to GP′ index. The RRM was used with complete phenotypic information for all individuals to estimate GEBV_ref_ over the gradient; and then independently repeated 10 times with the scenario CV-A to predict GEBV_pred_ for individuals in the validation set. GG_max_ was calculated as the mean of the top 5% of GEBV_ref_ and for each iteration GG_true_ was calculated as the mean of GEBV_ref_ associated to the top 5% individuals selected based on GEBV_pred_ for each GP′ values. GG_max_ and GG_true_ are centered and reduced.

#### Predictive performance over the CV scenarios

We considered an alternative cross-validation scenario (CV-B) (Fig. 2), to improve selection efficiency while preserving phenotyping effort with respect to CV-A. As in CV-A, 50% of the phenotypic data were used to constitute the T_set_ of the CV-B. The key difference between the two is due to a better distribution of phenotypic effort, both between individuals and between environments, in CV-B. This alternative distribution had a considerable impact on improving the predictive performance of the RRM, which increased from 0.25 for the CV-A to 0.59 for the CV-B (Fig. 8), with no increase in phenotyping effort. It should be noted that the CV-A scenario is a major challenge for RRM, as it imposes the prediction of entire trajectoriesforhalfofthepopulation.ThischallengeisrelaxedinCV-Bby including at least partial information for all individuals.

**Figure 8:**
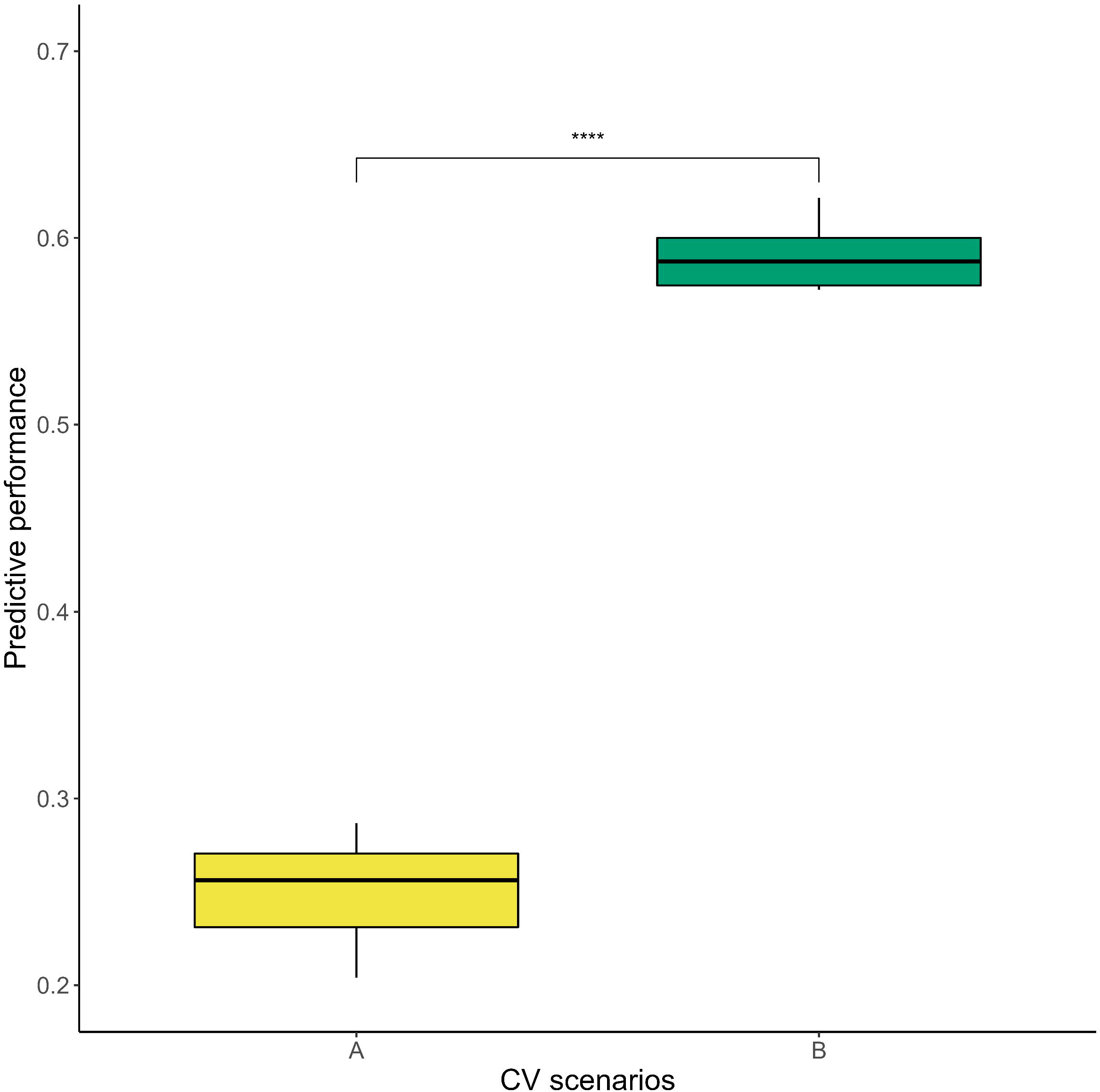
Predictive performance of the RRM according to the CV-A and CV-B scenarios. Boxplots indicates the Pearson correlation coefficient between observed and predicted RA values over the whole environmental gradient for 10 independent repetitions of the CV scenario. The significance between predictive performances was assessed by a Student’s t-test (****: p-value <1e10-4)

## Discussion

Deciding which genetic material should be planted now to form the forests of tomorrow is becoming increasingly challenging due to the rapidity of climate change (Thomas *et al*., 2004; Wiens, 2016). Using longitudinal tree-ring data and parallel environmental descriptors, we have successfully modeled genomic individual reaction norms based on random regression. This first example for forest trees provided consistent results for use in the maritime pine breeding program, but may inspire other programs in perennial species.

### Reaction norms in forest trees

Growth measurements at advanced age are generally used for the calculation of breeding values. Such measurements constitute highly integrative phenotypes that can be associated only with a global environmental site index. Using sites with contrasting indices has been a classic strategy to establish comparative trials for genetic x environment evaluation. In this sense, our two sites present strong contrast in terms of fertility and water table depth at the scale of the Landes massif, but even with their differences they are still part of the same breeding area (Jolivet *et al*., 2007). Wood cores give us access to phenotypic inter-annual variation and can be used to generate longitudinal annual growth data that can be associated with annual environmental variation. Our results showed indeed that the environmental variation between years was much greater than the one between sites (Fig. 5). Indeed Cir_22_ was associated with a mean environmental index GP′ of 68.2 for site 1 and 72.1 for site 2, whereas analysis based on ring measurements covered a larger index range (GP′ from 52.6 to 81.9). This much greater annual variation provides an opportunity to infer plasticity at individual level over a large environmental gradient.

In addition to longitudinal data collection, which can be operationally costly, there are other challenges that arise with these data. One is autocorrelation between repeated measurements on the same individual in a time series. Another, not least, is ontogenetic differences between phases of phenotype expression (Sanchez *et al*., 2013). Finally, a third challenge is the choice of a relevant environmental descriptor. Although we have not shown it for simplicity, we have performed a preliminary RRM for RA with a one-year lag in the climatic index in order to match RA of year *n* with the environmental index of year *n - 1*, and its results pointed to an absence of autocorrelated effect. As for the ontogeny challenge, we have ignored in our longitudinal data series the initial segments corresponding to the juvenile phase, keeping only the remaining adult phase for which the RA trend was generally flat, despite strong inter-annual oscillations (Fig. 5A).

The third challenge is probably the most difficult to address, the choice of a relevant environmental index (Li *et al*., 2017). This study was not designed to identify precisely the environmental factors most relevant to tree growth, but we defined two classes of biologically meaningful environmental indices that integrate the key components of temperature and water (Begum *et al*., 2013; Rathgeber *et al*., 2016). Both of them depend on the year and the site in which the ring was formed. The first class (aridity indices) is easy to obtain, since it only considers the climatic data (temperature and precipitation over the growing period) of the site and the year associated with to the rings under study. On the other hand, the second class (growth potential indices) requires more complex modeling, including for example the characterization of the daily water status of the trunk. A major difference between the two types of indices is the insensitivity of the former to the intra-annual distribution of precipitation and temperatures. Thus, similar annual aridity values (*DM* or *DM′*) may reflect different climatic realities over the course of the growing season, with temperatures and/or precipitation occurring at different periods and leading to differences in growth. Conversely, by considering the daily environmental status and tree physiology, the growth potential indices (*GP* and *GP′*) allowed a more detailed consideration of within-year environmental variation.

Finally, the predictive performance obtained with *DM*, *DM′*, *GP*, *GP′* (Fig. 4) confirms the relevance of the proposed environmental indices, but also suggest that they only partially capture the environmental factors influencing radial growth and the differences between individuals’ reactions. More specifically, the variability due to site is not fully described by the index, given the remaining high significance value of the corresponding fixed effect in RRM (Fig. 5B).

### Modelling reaction norms with RRM

Unlike univariate single-point analyses, which are easy to implement but do not integrate longitudinal phenotypic information, or multi-trait models, which can integrate it but are computationally demanding, RRM provides genetic estimates over the chosen continuous environmental gradient with reduced parametrization (Sun *et al*., 2017). The continuous trajectory of GEBV predicted by the RRM allows a position to be considered at any environmental level, whether it has actually been observed or not. The RRM can model highly complex curves using orthogonal base functions such as Legendre polynomials, which are widely used and described in the context of breeding (Schaeffer, 2004; Campbell *et al*., 2018; Marchal *et al*., 2019). Despite their great flexibility and computational advantages, Legendre polynomials may present numerical problems (Runge’s phenomenon) at the extremities for high-order fits (de Boor, 1978; Meyer and Kirkpatrick, 2005). In this study, the adjustment at the extremities of the environmental gradient was particularly important as the unfavorable extreme conditions are likely to increase in frequency in the future (Coumou and Rahmstorf, 2012; Spinoni *et al*., 2018). The use of low-order polynomials to model RA trajectories overcame this problem. The consistency and quality of the norms fitted with Legendre polynomials were verified by a comparison with norms fitted with B-spline functions, which are considered a more robust alternative to high-order polynomials in terms of extremum fitting, although less advantageous computationally (de Boor, 1978; Meyer and Kirkpatrick, 2005) (Kendall correlations between GEBV_ref_ estimated with Legendre polynomials and those estimatedwithB-splinesyieldingcoefficientsofupto0.95overtheentire gradient for the final RRM).

### Exploration of individual genetic trajectories

Random individual trajectories (Fig. 6) highlight the existence of plasticity for genetic values that can be targeted by breeders. It is not easy to discriminate between individual reaction norms that follow a trajectory close to the population average, given their high frequency, the fact that they present shallow slopes and mean GEBV_ref_ close to 0. However, individuals with potentially good growth along the entire gradient are much easier to discriminate from the rest, for which the proposed clustering allows simple and efficient visualization (cluster E), useful for selection purposes.

The distribution of individual GEBV varied between environmental levels, and those more favorable levels enhanced the expression of differences between trajectories relative to less favorable levels, which has already been observed in other biological models (Arnold *et al*., 2019). Genotype ranking was globally preserved over the trajectories for most of the gradient (van Eeuwijk *et al*., 2016). This trend was confirmed by strong correlations (supplementary Fig. S6) between the GEBV obtained with this RRM for radial growth and those obtained with the final circumference univariate analysis (final-point model). However, this correlation was weaker for unfavorable environments (environmental index below 59), in agreement with the reranking of genotypes observed for the individual trajectories at the most unfavorable end (Fig. 6). This precise and localized GxE interaction in our gradient, only possible thanks to the use of the RRM, should not be considered marginal or potentially negligible considering that it affects only one segment of the gradient. In fact, climate projections (supplementary Fig. S7) suggest that such unfavorable environments are likely to become much more common in the future. Even if the expected global level of aridity in 2075 remains close to current levels, according to our de Martonne calculation, aridity in 2100 will be much stronger, with a higher frequency of extreme events as predicted by other studies (Sillmann and Roeckner, 2008; Lehner *et al*., 2017). Our 15-year study period was already affected by a high global level of aridity and included extreme annual climates that may become frequent in the future. The environmental gradient used for the inference of reaction norms is therefore particularly relevant for identifying genotypes with better potential for growth in the unfavorable years to come.

When GxE interactions must be taken into account in selection decisions, a robust strategy would involve prioritizing the best adapted genotypes across the entire environmental gradient (Li *et al*., 2017), focusing on the notion of persistence. The definition of persistence may vary according to species and breeding aims (Gengler, 1996; Rocha *et al*., 2018), but it is generally defined as the capacity of a species to maintain a stable or high level of growth or production over time or in the face of different environmental conditions. For reactions norms, several ways of evaluating persistence and integrating the slope of trajectories in an operational breeding context have been proposed. For example, for feed conversion ratio in large white pigs, (Huynh-Tran *et al*., 2017) suggested combining the EBV estimated by the RRM with the coefficients of eigenvectors estimated from the eigenvalue decomposition of the covariance matrix of additive genetic effects. In their study, two summarized breeding values for each individual were sufficient to describe most of the variation in terms of mean genetic values (first dimension) and the slopes of EBV trajectories (second dimension), and could be used directly in selection. In another example in goat lactation, (Arnal *et al*., 2019) considered “the cumulative deviation in genetic contribution to yield relative to an average animal having the same (initial) yield” for the calculation of persistence-related EBV. Finally, (Peixoto *et al*., 2020) suggested the ranking of cotton genotypes on the basis of area under the reaction norm, the genotype with the highest norm being the most persistent. Another interesting approach would involve calculating the final GEBV for each individual as the mean of the GEBV for each environmental level weighted by the probability of occurrence of the environmental level in the future. Such a strategy would make use of the GxE interaction to maximize genetic gain for individuals performing in environmental conditions close to those predicted for the near future, while ensuring a certain level of resilience to environmental variation. Any of these proposals could be applied to our data. A possible advantage of the latter strategy could be to take more explicit account of future climate predictions, provided that they have some control over uncertainty.

### Reaction norm in a GS context

The use of genomic reaction norms to predict growth in unobserved environments is a good example of the potential benefits of GS approaches for traits that are complex to evaluate. Wood density profiles provide highly informative longitudinal data on tree growth over the years, but its acquisition via the coring process remains costly and time-consuming at breeding-program scale. This limitation has motivated one of our alternative cross-validation scenarios (CV- B), with a more homogeneous distribution of phenotypic effort, resulting in a training population involving all individuals, 25% of which contribute full time series and the remaining 75% only partial 5-year series. Indeed, relative to our baseline scenario (CV-A), which aimed to predict the full trajectories of 50% of individuals, the CV-B scenario achieved a much higher level of predictive performance (0.59), demonstrating that the allocation of phenotyping effort to constitute the training population is a key optimization to consider. The scenario CV-B would reflect the use of a high-throughput phenotyping tool usable on a large number of individuals at the cost of a smaller number of rings scanned per tree, which is basically what a resistograph does (Bouffier *et al*., 2008). Resistograph measures the resistance of the wood to penetration with a needle and can estimate RA efficiently for the rings closest to the bark, i.e. the last five rings formed (personal communication). These measurements provide only partial information about plasticity, but when applied to the whole population, they have the advantage of providing information complementary to that obtained by coring. Overall, less phenotyping effort is required, but the benefits are substantial.

The genetic component of reaction norms, the one of greatest interest to breeders, was estimated by integrating pedigree or genomic information in the RRM. Genomic-based RRM had a significantly better predictive performance (with the *GP′* index) than pedigree-based RRM, suggesting that refining the coefficients of relationships between individuals through their molecular characterization with SNPs results in the generation of more suitable models (Gamal El-Dien *et al*., 2016; Bouvet *et al*., 2016). The pedigree information tended towards a systematic overestimation of pairing coefficients relative to the genomic information (Fig. 3). However, some rare pairs of individuals appeared to be much more related on the basis of genomics than on the basis of pedigree, suggesting that, in some cases, the pedigree may be incomplete, or may contain errors, despite the correction and recovery steps (Tan *et al*., 2017; Li *et al*., 2019). The use of genomic data for genomic evaluation is often proposed for forest trees (Grattapaglia and Resende, 2011; Lebedev *et al*., 2020), but first GS studies for maritime pine (Isik *et al*., 2016; Bartholomé *et al*., 2016) highlighted the difficulty of demonstrating a superiority of genomic models over pedigree-based models. In this study, we provide some arguments to go beyond these limitations in the application of the genomic prediction model. The RRM takes greater advantage of genomic information to predict individual trajectories than pedigree information. Indeed, in a context of intense climate change, the importance of integrating environmental information into genetic evaluation may fully justify the additional cost of genotyping (Isik, 2014).

## Acknowledgments

This work was supported by the European Union’s Horizon 2020 Research and Innovation Programme Project under grant agreement n°773383 (B4EST). VP was awarded a doctoral fellowship (N°2020-CK-126) from Ecole Nationale Supérieure Des Sciences Agronomiques de Bordeaux-Aquitaine, 1 cours du Général de Gaulle, CS 40201 33175 Gradignan Cedex. The authors would like to thank GIS “Groupe Pin Maritime du Futur” and INRAE - UEFP (https://doi.org/10.15454/1.5483264699193726E12) for the installation of the studied sites, the management of the sites, the help to collect data (circumference measurements) and biological material (needles and increment cores). Authors are thankful to Frederic Lagane (PHENOBOIS Platform) for the cutting and the radiography of the increment cores. Authors thank the Institute of Biosciences and BioResources – Italian National Council of Research (IBBR/CNR), especially Giovanni Giuseppe Vendramin and Sara Pinosio for performing most of DNA extraction and quality monitoring. Part of the experiments (DNA extraction, quantification and manipulation) were also performed at the PGTB (doi:10.15454/1.5572396583599417E12), with the help of Christophe Boury and Céline Lalanne. Raphaël Segura provided soil and climate characterization for the two studied sites. Authors are also thankful to Christophe Plomion for his help during the conceptualization of this study.

## Author contributions

LB and LS: conceptualization, supervision and validation

VP, AB, LB and LS: methodology

VP: data curation, formal analysis, visualization, writing – original draft preparation

VP and AB: software

AB, LB and LS: writing – Review & Editing

## Data Archiving

The data underlying this article are accessible via the private following link (Data INRAE): https://entrepot.recherche.data.gouv.fr/privateurl.xhtml?token=15f2101e-ebb8-4b7c-838b-9703090cfec4

The corresponding DOI is https://doi.org/10.57745/NUTK1I (Papin, Victor; Bosc, Alexandre; Sanchez-Rodriguez, Leopoldo; Bouffier, Laurent, 2023)

The data will be made public and accessible to all once the article has been accepted.

## Research Ethics Statement

Not applicable.

