## Supplementary_information for "Integrating environmental gradients into breeding: application of genomic reactions norms in a perennial species"

#### Supplementary Tables

Table S1

Table S2

#### Supplementary Figures

Figure S1

Figure S2

Figure S3

Figure S4

Figure S5

Figure S6

Figure S7

#### Supplementary Methods

Method.S1

Method.S2

**Table S1: Soil and climate characterization for Site 1 and Site 2**

| Site | Soil characterization[1] | | | | Climate characterization[2] | | | | | | | |
|---|---|---|---|---|---|---|---|---|---|---|---|---|
| | *Organic matter (g/kg of soil)* | *Water table depth (m)* | | | *Cumulative annual rainfall (mm)* | | | | *Mean annual temperature (°C)* | | | |
| | *Mean* | *Min* | *Mean* | *Max* | *2015* | *2016* | *2017* | *2018* | *2015* | *2016* | *2017* | *2018* |
| Site 1 | 37.3 | 0 | 0.9 | 1.8 | 407 | 492 | 582 | 503 | 16.2 | 15.5 | 16.4 | 16.4 |
| Site 2 | 20.5 | 5.8 | 6.9 | 7.8 | 511 | 522 | 647 | 585 | 16.4 | 15.9 | 16.5 | 16.4 |

[1] Soil organic matter content was determined by an analysis of the first 80cm of soil performed in 2015; water table depths were recorded from 2015 to 2016 with soil humidity probes
[2] Climate data are from weather stations located on each site

**Table S2: Number of rings available for POP after filtering according to the year**

| Year | *1998* | *1999* | *2000* | *2001* | *2002* | *2003 to 2017* | *2018* | *2019* |
|---|---|---|---|---|---|---|---|---|
| Number of rings available for POP | 3 | 203 | 530 | 606 | 624 | 628 | 627 | 625 |

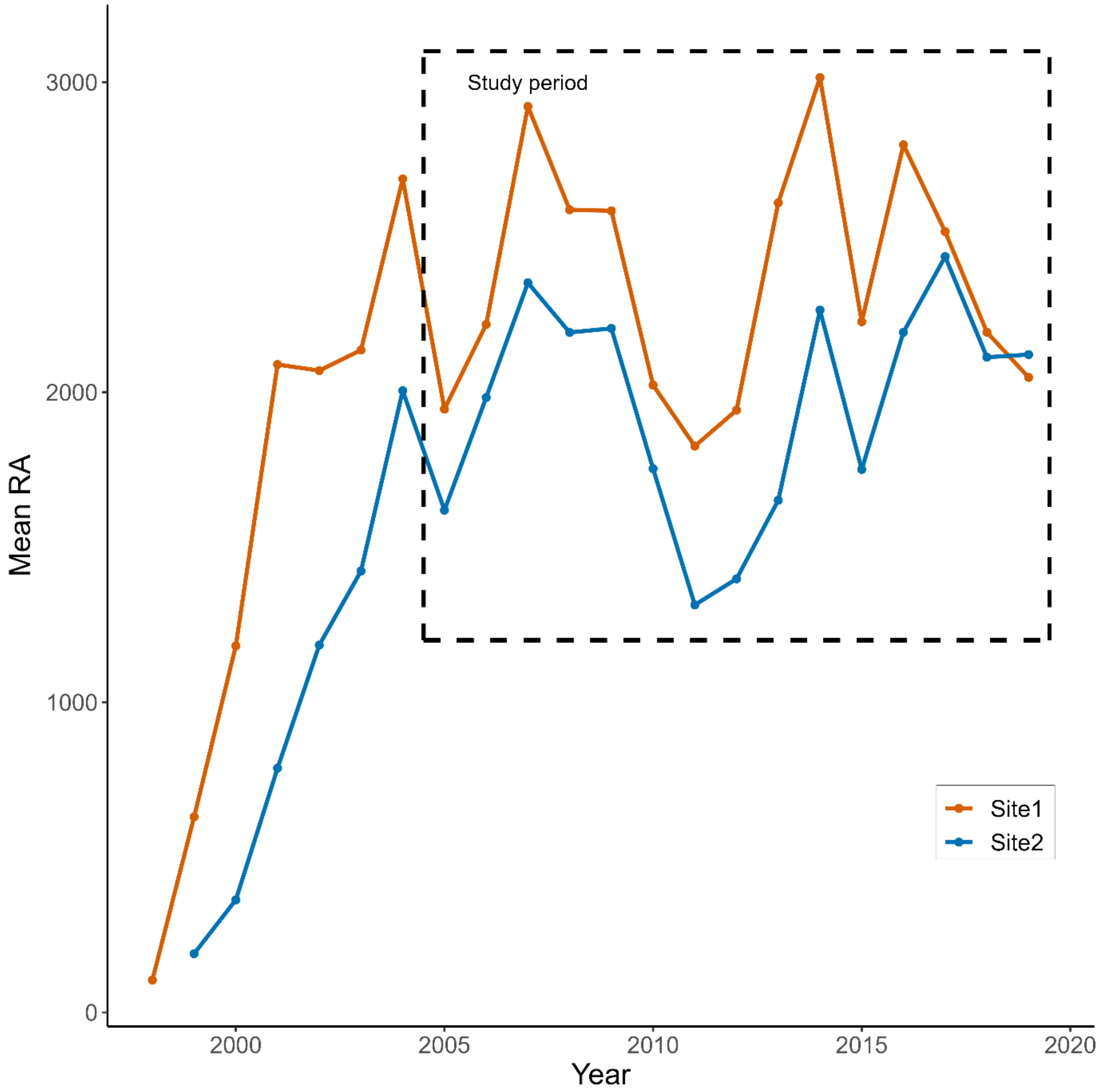


**Figure S1: Evolution of mean RA over the years for individuals of POP.** *The orange and blue lines represent the average trajectories of the 303 individuals of Site1 and the 325 individuals of Site2, respectively. Our study period from 2005 to 2019 is framed in a dotted box.*

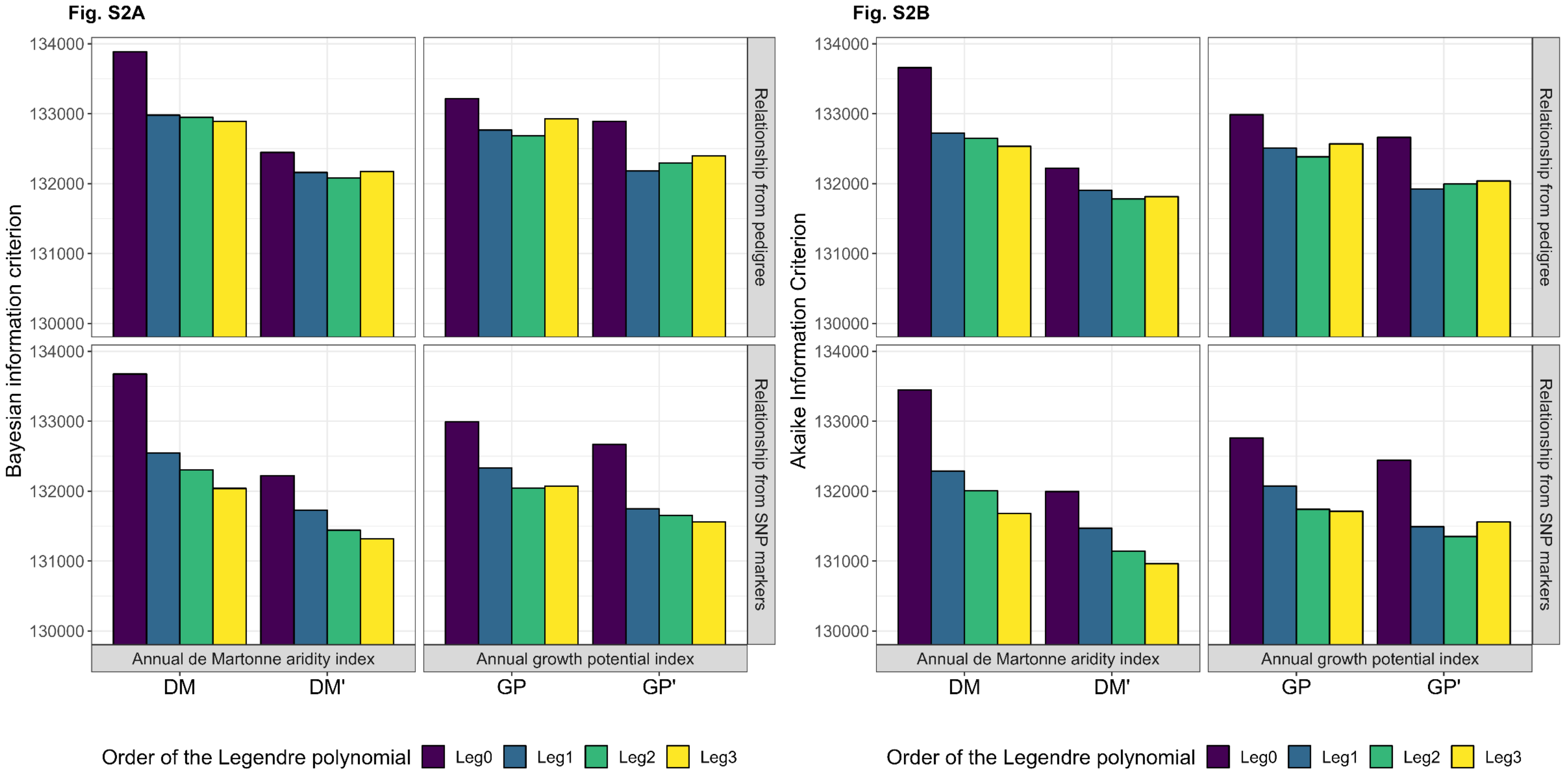


**Figure S2: Comparison of Bayesian information criterion (BIC) (Fig. S2A) and Aikaike information criterion (AIC) (Fig. S2B) for RRM with different orders of Legendre polynomials.** *Order 2 RRMs seem more appropriated in our study to model the relatively simple trajectories of RA. Compared to order 3 RRMs, order 2 RRMs display very similar shapes of reaction norms and allow a drastic reduction on computational demand, with a marginal loss of goodness of fit according to AIC and BIC when appearing as second the second best fit.*

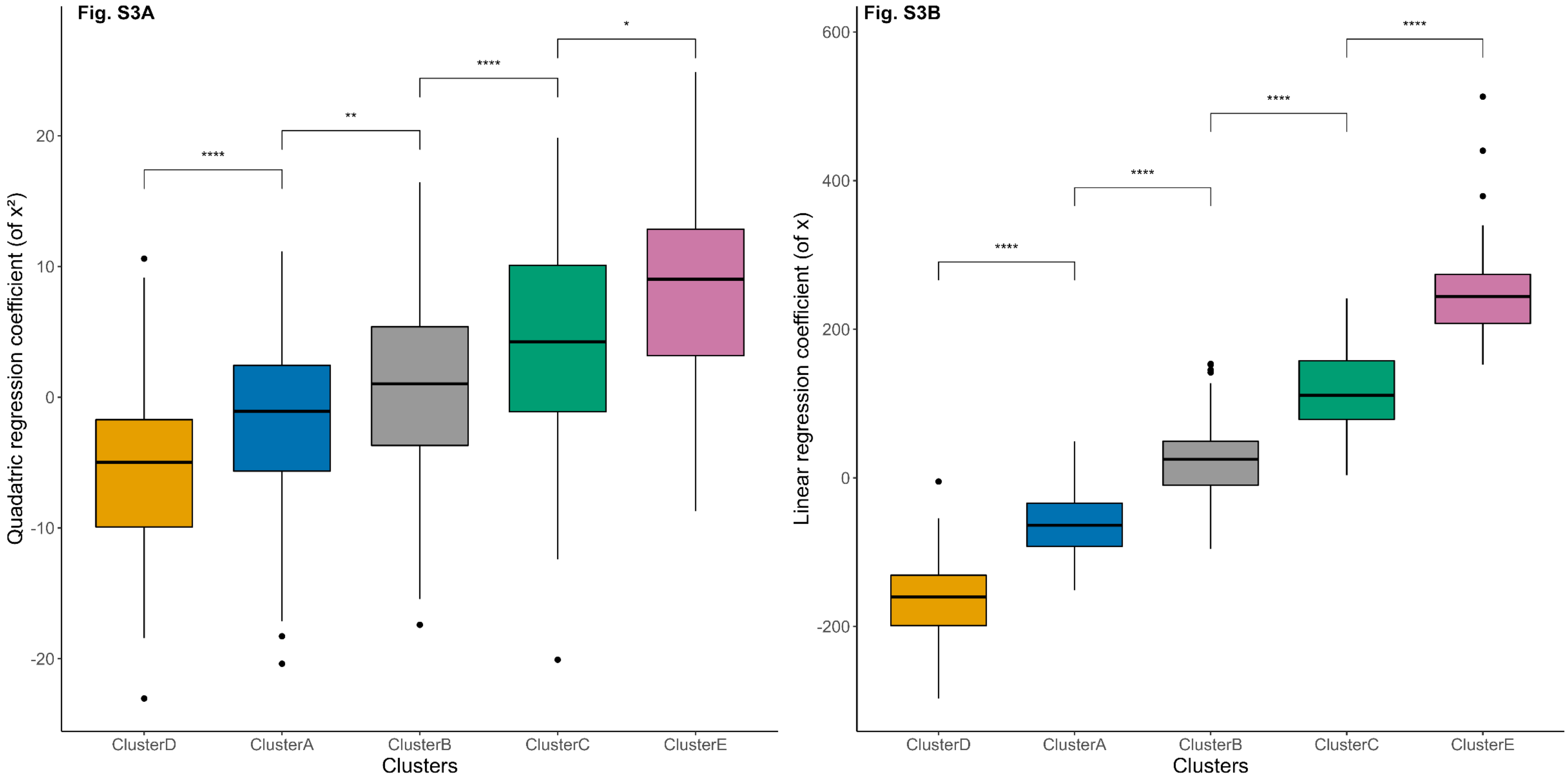


**Figure S3**: **Quadratic and linear regression coefficients of the trajectories estimated by the RRM depending on the cluster.** *The differences between boxplots were assessed by a Student's t-test and the significance level associated is indicated above boxplots (*: p-value<0.05, **: p-value<0.01, ****: p-value<0.0001)*

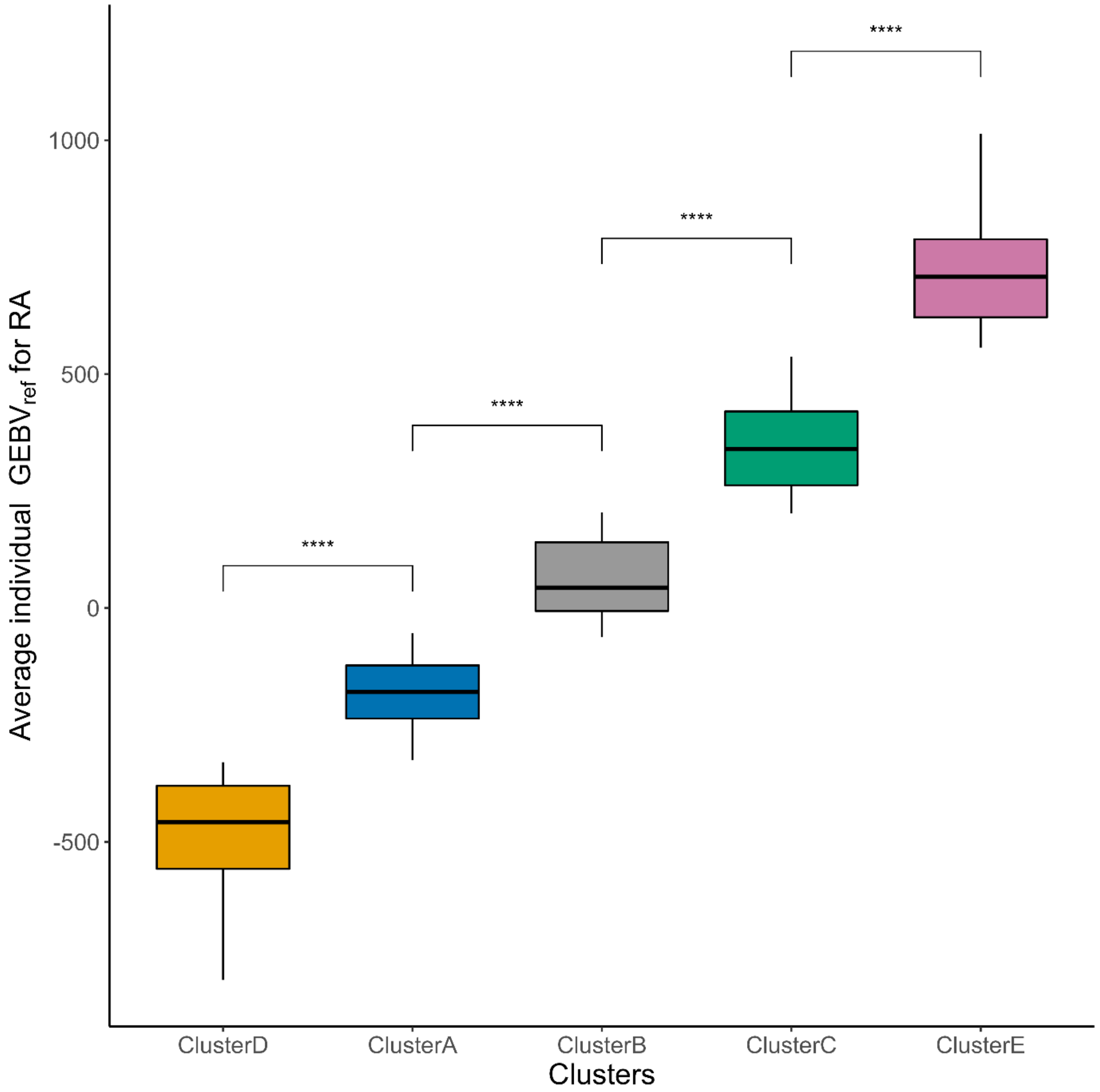


**Figure S4: Mean $GEBV_{ref}$ per individual estimated by the RRM depending on the cluster. The differences between boxplots were assessed by a Student's t-test and the significance level associated is indicated above boxplots (****: p-value<0.0001)**

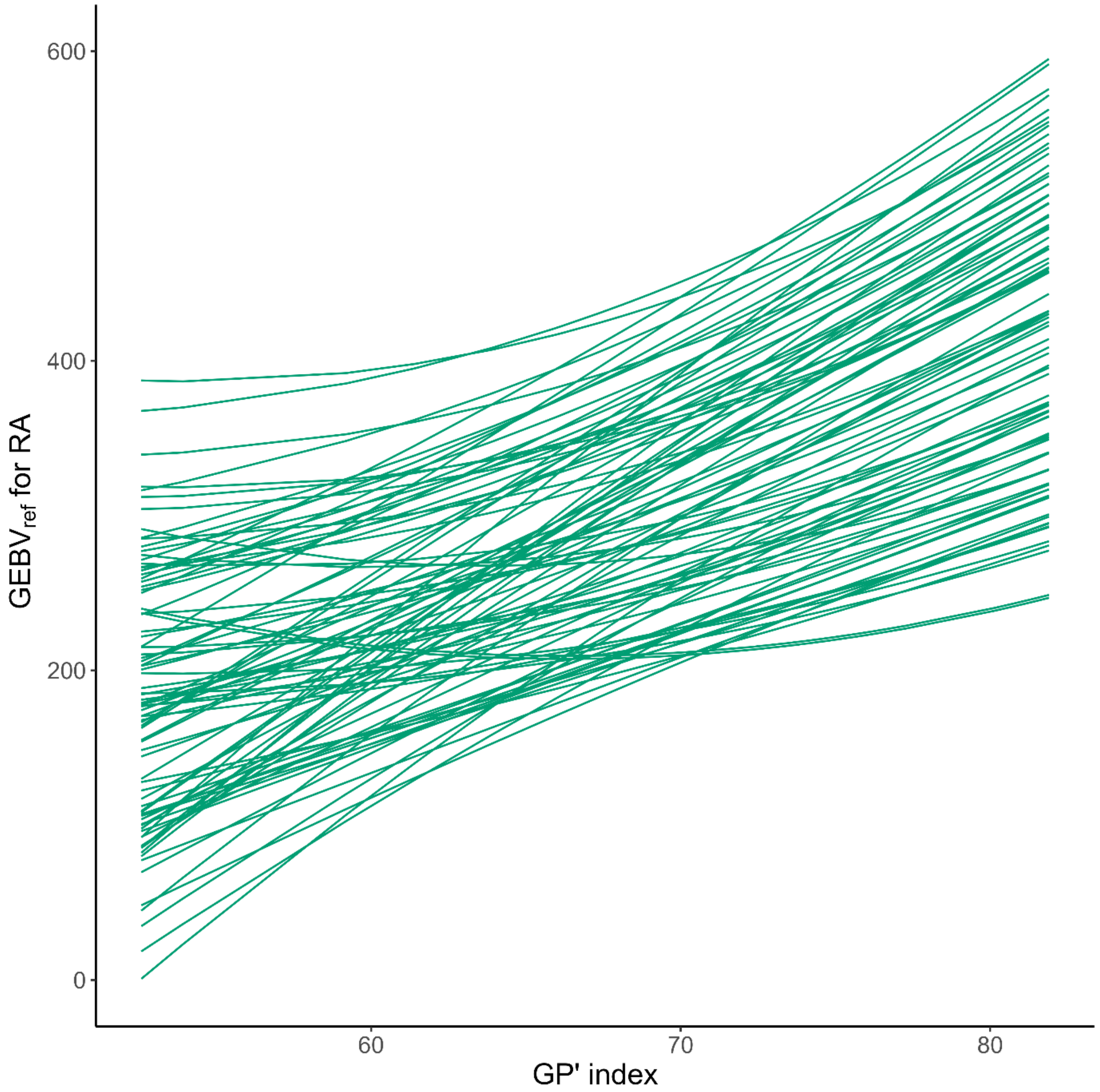


**Figure S5: Trajectories of individual GEBV$_{ref}$ as a function of annual GP' index estimated by the RRM, only for individuals from cluster C (18.6%)**

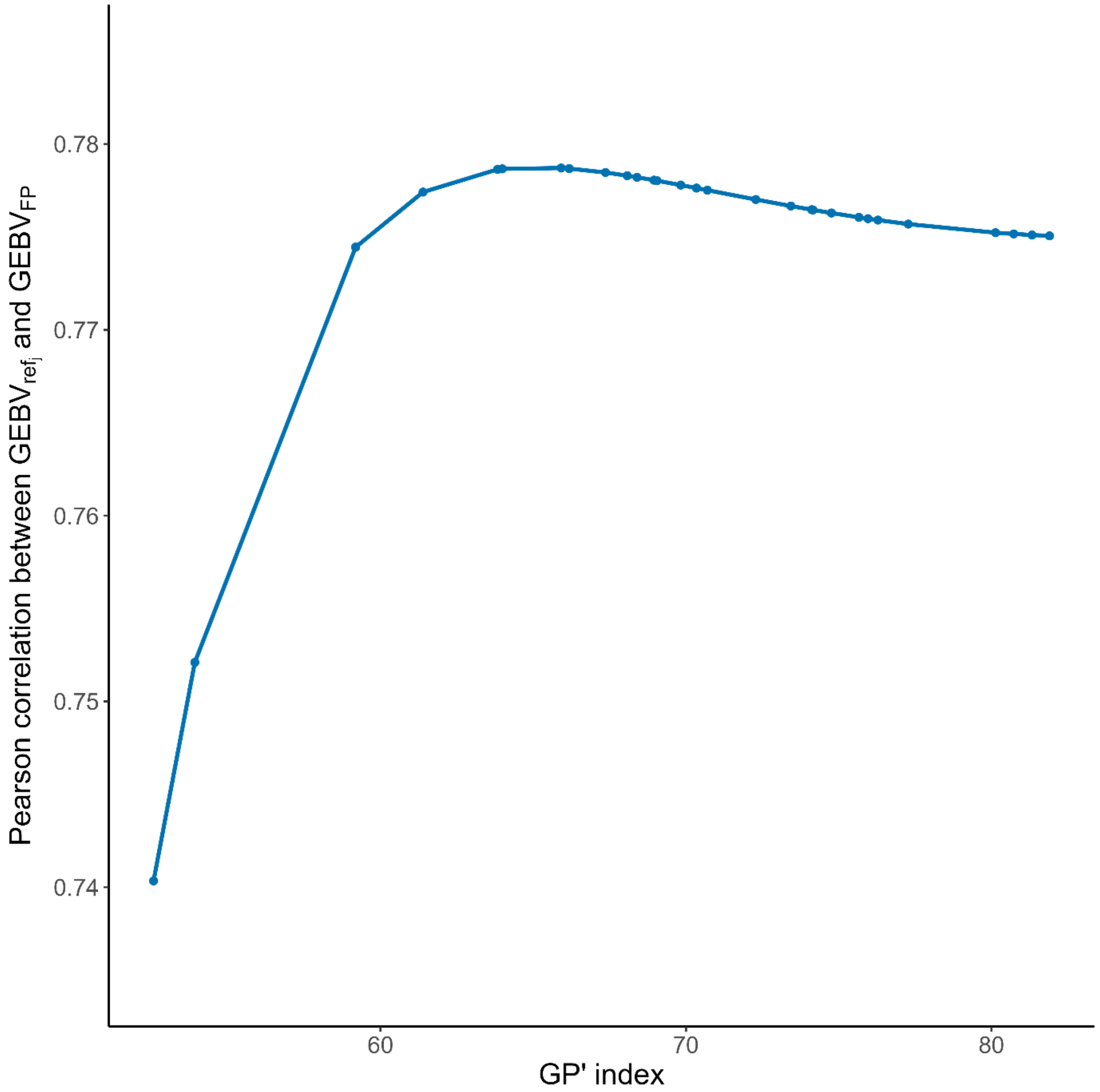


**Figure S6: Pearson correlation coefficients between genomic estimated breeding values obtained from a final-point univariate model ($GEBV_{FP}$) and genomic estimated breeding values obtained from a RRM at each GP' level j ($GEBV_{ref\_j}$)**

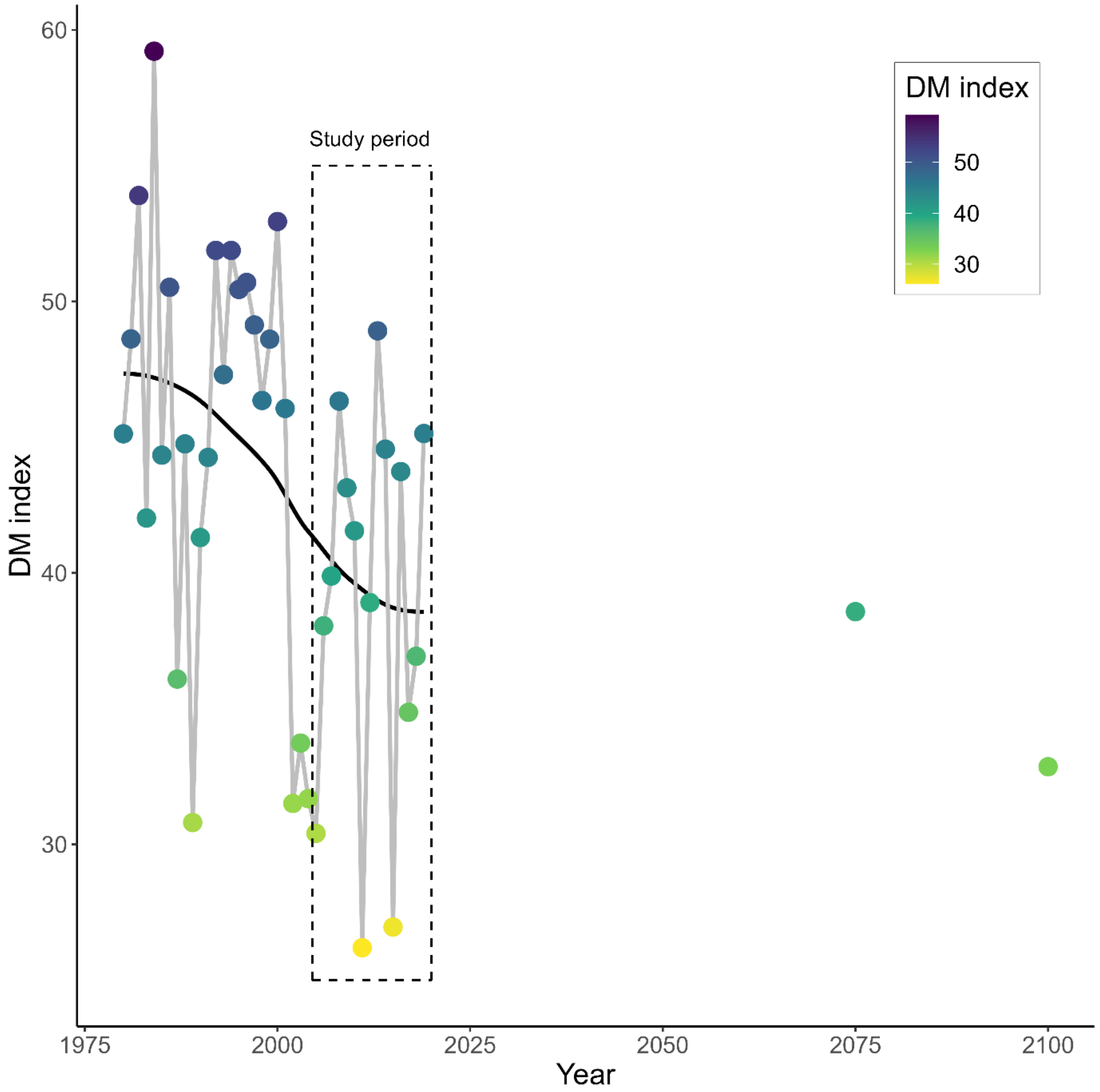


**Figure S7**: **Evolution of DM index from 1980 to 2019 and prediction of this index for medium and long term horizon**. *Evolution from 1980 to 2019 was calculated with historical data from Meteo France weather station close to Site1. Predictions positioned in 2056 and 2086 were calculated using weather predictions available on drias-climat.fr considering the RCP8.5 scenario (Scenario without any climatic politic) for period from 2041 to 2070 (medium horizon) and from 2071 to 2100 (long term horizon). Our study period from 2005 to 2019 is framed in a dotted box.*

### Method S1. Pedigree recovery for POP

***Preliminary stage***

The 25 known female parents and the 85 potential male parents (corresponding to two pollen mixtures from 42 and 43 male parents) of POP are grafted in clonal archives and needles were sampled for DNA extraction. Following the same procedure as for POP, an additional genomic relationship matrix (noted $G_{all}$) was computed with the 628 individuals of POP and the 85 parents that successfully passed the quality controls. $G_{all}$ $(713\ x\ 3832)$ was used only for pedigree recovery, as described below.

For the small proportion of parents (1/25 mothers and 24/85 fathers) that did not pass the quality controls, genotyping information was available for a limited number of SNP markers from a previous study (Vidal *et al.*, 2017).

***Criteria for pedigree recovery***

A subset of 161 highly discriminant SNPs (LD-$r^2$<0.1 and MAF>0.4 in POP) was chosen from the 4TREE array and used for pedigree recovery. The 25 parents not successfully genotyped with the 4TREE array were only characterized for only 21/161 SNPs (available from Vidal et al., 2017). For the female and male parents, the two most likely candidates were identified with Cervus 3.0 (Marshall *et al.*, 1998; Kalinowski *et al.*, 2007), as described by Vidal et al. (2017). The candidate considered most likely was examined first, followed by the second most likely candidate if necessary. For validation, a candidate parent had to have (i) fewer than two mismatches with the descendant and (ii) a high level of relatedness ($0.5 \pm 20\%$) to the descendant (relatedness coefficient extracted from $G_{all}$). In the specific case in which information for only 21 SNPs was available for the candidate parent, criterion (ii) was replaced with the delta score criterion (estimated with Cervus 3.0 and described by Vidal et al., (2017).

***References Supplementary Method S1***

**Kalinowski ST, Taper ML, Marshall TC**. 2007. Revising how the computer program cervus accommodates genotyping error increases success in paternity assignment. Molecular Ecology **16**, 1099–1106.

**Marshall TC, Slate J, Kruuk LEB, Pemberton JM**. 1998. Statistical confidence for likelihood-based paternity inference in natural populations. Molecular Ecology **7**, 639–655.

**Vidal M, Plomion C, Raffin A, Harvengt L, Bouffier L**. 2017. Forward selection in a maritime pine polycross progeny trial using pedigree reconstruction. Annals of Forest Science **74**, 21.

### Method S2 – Environmental indices

#### A. Modified de Martonne aridity index

The modified version of the de Martonne index proposed in this study is based on the formula suggested by Botzan et al. (1998):

$$DM'_{y,z} = \alpha\, DM_{y,z} + (1-\alpha)DM_{(y-1),z}$$

Where $DM'_{y,z}$ is the modified de Martonne index for the year $y$ and site $z$; $DM_{y,z}$ and $DM_{(y-1),z}$are respectively the de Martonne index for the year $y$ and $y-1$; and $\alpha$ is a coefficient defined as below:

$$r = \frac{DM_{y,z} - DM_{(y-1),z}}{DM_{y,z}} \quad , \quad if \begin{cases} r > 0.5 \\ r = 0.2\ to\ 0.5 \\ r < 0.2 \end{cases} \quad then \begin{cases} \alpha = 0.75 \\ \alpha = 0.90 \\ \alpha = 1.00 \end{cases}$$

We tested different threshold values for $r$ classes $\{(0.5-0.2),(0.4-0.2)\}$ and for $\alpha$ classes $\{(0.75-0.90-1.00),(0.50-0.75-1.00),(0.25-0.50-1.00),(0.00-0.25-1.00)\}$. Each time, the gradient of annual de Martonne indices obtained was used in a RRM and the quality of the model was assessed by a cross-validation routine (CV-A). The best model quality was obtained with the annual modified de Martonne index gradient calculated on the sliding window of 30 days with $r$ class = $\{(0.4-0.2)\}$ and $\alpha$ class = $\{(0.25-0.50-1.00)\}$. The final formula used is thus:

$$DM'_{y,z} = \alpha\, DM_{y,z} + (1-\alpha)DM_{(y-1),z}$$ , with:

$$r = \frac{DM_{y,z} - DM_{(y-1),z}}{DM_{y,z}} \quad , \quad if \begin{cases} r > \mathbf{0.4} \\ r = \mathbf{0.2}\ to\ \mathbf{0.4} \\ r < \mathbf{0.2} \end{cases} \quad then \begin{cases} \alpha = \mathbf{0.25} \\ \alpha = \mathbf{0.50} \\ \alpha = \mathbf{1.00} \end{cases}$$

#### B. Inputs and details for GO+ 3.0 model

A growth potential index (GP) was calculated for each year $y$ within each site $z$, based on mean trunk water potential ($\varphi_{trunk}$) and temperature ($T_a$) estimated daily by the GO+ v3.0 model (Moreaux *et al.*, 2020):

$$GP_{y,z} = \sum_{i=1}^{365} GP_{\varphi_{tronc_i}} . GP_{T_{a_i}}$$

where $GP_{\varphi_{trunk}}$ and $GP_{T_a}$ are the growth potential components linked to $\varphi_{trunk}$ and $T_a$, respectively. $GP_{\varphi_{trunk}}$ and $GP_{T_a}$ are obtained for each day $i$ with the following response functions:

$$GP_{\varphi_{trunk_i}} = \frac{1}{1 + e^{-\lambda\,(\varphi_{trunk}+c)}} \quad and \quad GP_{T_{a_i}} = Q_{10}^{\frac{T_a - T_{ref}}{10}}$$

with the following parameters for the sigmoid function: $\lambda\ (slope) = 10$ and $c = \frac{\max(\varphi_{trunk}) - \min(\varphi_{trunk})}{2}$ (an additional parameter centering the sigmoid on our range of $\varphi_{trunk}$ values), and the following values for the Q10 function: $Q_{10} = 10$ and $T_{ref} = 29.9\ (°C)$ (maximum temperature $T_a$ over our study period 2005-2019).

The GO+ model was used with maritime pine species parameters (default). Soil parameters used for Site1 and Site2 were respectively typical of wet Landes and dry Landes with low fertility. Real climatic and silvicultural data were added.

The GO+ model produced stand-level average values of microclimate temperature ($T_a$) root water potential ($\varphi_{roots}$) and canopy water potential ($\varphi_{canopy}$), at a daily scale over the period 2005-2019. These last two parameters were combined to estimate a trunk water potential at 1.30m ($\varphi_{trunk}$):

$$\varphi_{trunk} = \varphi_{roots} - \frac{1.3 + 0.7}{Ht_{mean} + 0.7} \left(\frac{\varphi_{roots}}{\varphi_{canpy}}\right)$$

With $Ht_{mean}$ the average stand height for the given day.

Note that the GO+ version used does not incorporate growth data from our stands. The independence between environmental variables and phenotypic trait used in the RRM is therefore guaranteed.

#### C. Modified Growth Potential index

From the initial GP calculation procedure, different values of slopes $\alpha = \{20, 10, 5, 2\}$ and center of the curve $c = \{c_{ref}$-50%, $c_{ref}$-25%, $c_{ref}$, $c_{ref}$+25%, $c_{ref}$+50%} were tested for the sigmoid response function to $\varphi_{trunk}$, as well as different values of $Q_{10}$={2,3,4} were tested for the response function to temperature $T_a$. To get the final growth potential for one year, different types of annual integration of $GP_{\varphi_{tronc_i}} . GP_{T_{a_i}}$ were tested (sum of daily values over a year; average over a sliding window of 2, 5 or 10 days before summing over a year; assignment of the minimum obtained over a sliding window of 2, 5 or 10 days before summing over a year). In addition, modifications to the annual GP values to take into account the impact of previous year were applied in the same way as for the modified de Martonne index. The best model quality (i.e. best predictive performance for CV-A) was obtained using an annual environmental gradient calculated with the initial procedure and initial values (sigmoid function response to $\varphi_{trunk}$ with $\alpha = 10$ and $c = c_{ref}$; Q10 function response to $T_a$ with $Q_{10} = 2$) but using the minimum value of $GP_{\varphi_{tronc_i}} . GP_{T_{a_i}}$ in a sliding window of 10 days before for the annual integration and considering the impact of previous year (modifications with $r\ classes\ \{(0.2 - 0.4)\}\ and\ \alpha\ \{\ (0.25 - 0.50 - 1.00)\}$).

The final formula for the GP' index is thus: $GP'_{y,z} = \alpha\ GP_{y,z} + (1 - \alpha) GP_{(y-1),z}$ , with: (1) and (2)

$$(1): GP_{y,z} = \sum_{i=1}^{365} min\ \left\{GP_{\varphi_{tronc_{i-10}}} . GP_{T_{a_{i-10}}}, \ldots, GP_{\varphi_{tronc_i}} . GP_{T_{a_i}}\right\}$$

When $i \in [1:10]$, $GP_{\varphi_{tronc_{i-10}}} . GP_{T_{a_{i-10}}}$ are from the previous year (last days of December)

$$(2): r = \frac{GP_{y,z} - GP_{(y-1),z}}{GP_{y,z}}\ ,\ \ if \begin{cases} r > \mathbf{0.4} \\ r = \mathbf{0.2}\ to\ \mathbf{0.4} \\ r < \mathbf{0.2} \end{cases} then \begin{cases} \alpha = \mathbf{0.25} \\ \alpha = \mathbf{0.50} \\ \alpha = \mathbf{1.00} \end{cases}$$

***References Supplementary Method S2***

**Botzan TM, Mariño MA, Necula AI**. 1998. Modified de Martonne aridity index: application to the Napa Basin, California. Physical Geography **19**, 55–70.

**Moreaux V, Martel S, Bosc A, *et al.*** 2020. Energy, water and carbon exchanges in managed forest ecosystems: description, sensitivity analysis and evaluation of the INRAE GO+ model, version 3.0. Geoscientific Model Development **13**, 5973–6009.
